# Pangenome-based human genome analysis improves trait association and genomic prediction

**DOI:** 10.64898/2026.07.01.735728

**Authors:** Shuangjia Lu, Wen-Wei Liao, Marianne K. DeGorter, Page C. Goddard, Jana Ebler, Tsung-Yu Lu, Human Pangenome Reference Consortium, Mark J. P. Chaisson, Tobias Marschall, Stephen B. Montgomery, Nathan O. Stitziel, Ira M. Hall

## Abstract

The Human Pangenome Reference Consortium has generated 462 open-access reference genomes and a variation graph that represents differences among them, providing a substrate for pangenome-based analysis methods that overcome the longstanding limitation of comparing all genomic data to a single linear reference. A key unresolved question is the extent to which these approaches can improve trait mapping. We investigate this using the genetics of gene expression variation as a model. We developed a graph-based method (EdgeDepth) for associating sequence variation with traits using short-read genome sequencing data, and show that it captures complex forms of genetic variation missed by other methods. We evaluated trait mapping performance using 430 samples with deep RNA-seq data, and found that pangenomic methods enable the detection of expression quantitative trait loci involving multiallelic indels and structural variants, leading to increased power at a subset of genes. These include 812 genes (7.9% of total) with ≥20% improvement in statistical significance relative to the 1000 Genomes Project callset, and 185 (1.8%) with a 50% improvement, 10 of which are candidates to explain prior GWAS results. Notably, these analyses implicate *GBAP1* pseudogene copy number as a causal factor in Crohn’s disease, likely via miRNA-mediated regulation of *GBA1*, which explains prior GWAS results based on flanking SNPs. The inclusion of pangenome-specific variation also improved the performance of gene expression prediction models, with median variance explained increasing from 10.1% to 12.5%, and 14.6% of genes showing significant improvement (Δr^2^>0.05). Taken together, these results suggest that integration of pangenomic methods into human genetic studies will improve trait association and genomic prediction at a meaningful subset of genes.

## Introduction

The new data resource released by the Human Pangenome Reference Consortium (HPRC) includes 462 high quality genome assemblies, pangenome graphs created by whole-genome multiple sequence alignment, and a comprehensive variant catalog that includes complex and repetitive variant types that have historically been recalcitrant to analysis using short-read data and linear references^1^. This new resource (HPRC2) builds upon our earlier release (HPRC1) that included 94 haplotypes^2^. These resources are facilitating the development of a new generation of pangenome-based bioinformatics tools that aim to leverage allelic information from diverse haplotype sequences to improve genome analysis in human genetic studies and clinical sequencing efforts. These include relatively mature tools for read alignment (e.g., Giraffe^3^) and variant calling (e.g., PanGenie^4^), plus a host of other methods at various stages development. A key promise of these methods is that they will allow many new genetic variants to be accurately genotyped and tested for association in human disease studies even when only short-read whole-genome sequencing (WGS) are available (as is typically the case).

However, the extent to which pangenome variant analysis tools will improve trait mapping remains an open question. There is clear evidence that long-read sequencing and whole-genome assembly methods are capable of identifying many novel variants that are not detectable by aligning short-read WGS data to a linear reference genome. For example, current large-scale structural variant (SV) callsets based on short-read data typically have ∼5,000-10,000 SVs per sample^5–7^, whereas long-read genome assembly methods identify ∼25,000-30,000 SVs^2,8–10^. Once these missing variants have been mapped to base-pair resolution using long-read data and their precise structures encoded in pangenome references, they can often be genotyped using short-reads^2,8^. What is far less clear is the extent to which these missing variants are relevant to disease given that most are found in repetitive regions of the genomes that are relatively unconserved and depleted for known functional elements, and thus are less likely to be functionally relevant. There are also open questions about the best way of detecting and testing these variants, many of which are multiallelic SVs and variable number tandem repeats (VNTRs) with complex structures and repetitive breakpoints that are extremely challenging to measure accurately. Understanding the functional significance of these variant classes and the best approaches for measuring them is important because the reanalysis of population-scale short-read WGS datasets with pangenomic methods will be a costly and laborious task that should be guided by the empirical value of these approaches.

In this study we investigate the utility of pangenomic approaches by developing EdgeDepth, a new method that leverages pangenome graphs for trait association, and by comparing its performance (in combination with other pangenomic methods) to the current gold standard for short-read variant mapping, the 1000 Genomes Project (1KG) callset^7^. The 1KG callset relied on GATK^11^ for small variants plus a multi-group, multi-algorithm SV discovery effort that included many difficult-to-detect variants, including multiallelic copy number variants (CNVs) and novel insertions of sequence not in the GRCh38 reference genome. As such, although reference-based variant calling methods have improved to a minor degree in the last few years, the published 1KG callset is a reasonable approximation of the best-case scenario for short-read WGS using the standard linear reference (GRCh38).

We use expression quantitative trait locus (eQTL) mapping and cis-heritability of expression variation as a model for what pangenomic approaches might be expected to contribute to the analysis of complex human traits. Studying genetically regulated gene expression is informative for complex trait genetics because both GWAS hits and eQTLs are typically caused by noncoding variants that affect gene expression, the spectrum of causal functional variants is expected to be similar – at least in terms of variant types and detectability – and in both cases the identification and fine-mapping of association signals is dependent on sensitive and accurate variant calling. Thus, gains demonstrated through eQTL mapping are likely to generalize to larger-scale studies of more complex traits.

## Results

### The EdgeDepth method

We first developed EdgeDepth, a new method for direct association of pangenome graph features with traits. The EdgeDepth analysis pipeline includes (**Fig. 1**): (1) alignment of WGS reads to a pangenome graph, in this study using short-read data, the HPRC2 graph generated by the Minigraph-Cactus pipeline^12^, and the Giraffe aligner^3,13^; (2) measurement of aligned read-depth at graph edges; (3) cross-sample read-depth normalization; (4) filtering redundant edges so that each variant allele is represented by a single edge, retaining the edge with the most read-depth support; (5) selection of high confidence variable edges that have reliable alignment depth and show variability across samples (in this case present in ≥10 samples); (6) refinement of read-depth information at biallelic variants by combining information from the 4 constituent edges into a single “allele balance” value; and (7) performing trait association using the quantitative allele balance and edge depth values as inputs. A key advantage of this method is it is simple and, by testing the copy number of graph edges, it allows any sequence feature to be associated with a trait, without requiring accurate knowledge of allelic structure. A second advantage is that, unlike other pangenomic approaches that use k-mer counts^4,14^, EdgeDepth uses full read alignment, which is expected to be more accurate in repetitive sequences.

**Figure 1.**
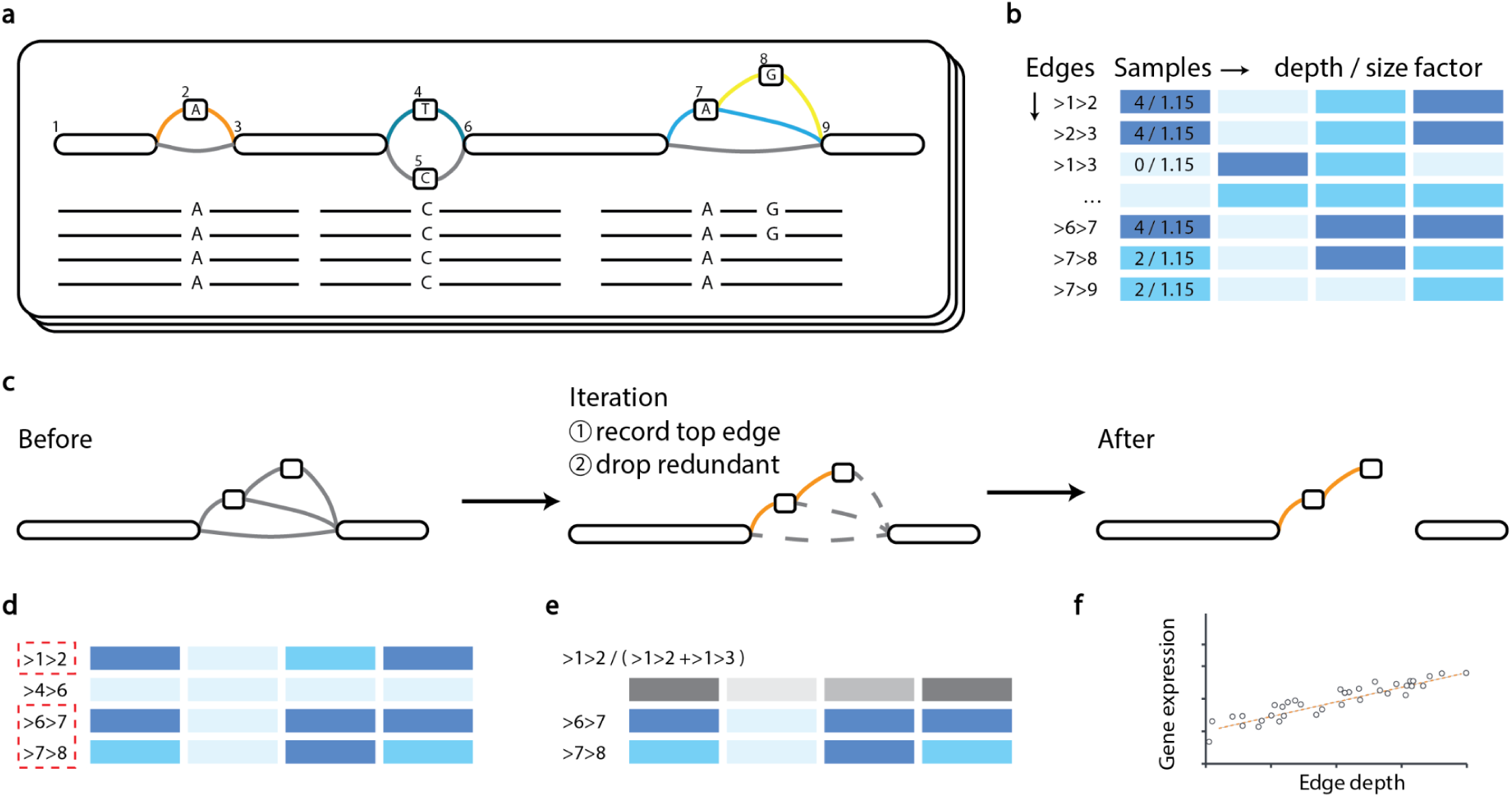
Overview of the EdgeDepth pipeline. EdgeDepth quantifies variants using graph-based read alignment and tests them for association with traits. Throughout, an edge connecting node x to node y is denoted as >x>y, where > indicates that the following node is traversed in the forward orientation and < indicates traversal in the reverse-complement orientation. In heatmaps, columns represent samples, rows represent edges, and color intensity encodes the edge depth value. **(a)** Step 1: align short reads from WGS data of each sample to a pangenome reference graph. Numbered segments are nodes, and colored arcs are edges carrying alternative alleles, with the WGS reads shown below. **(b)** Steps 2-3: quantify read support for each graph edge in each sample, then normalize edge depth by a per-sample size factor to make depths comparable across samples. **(c)** Step 4: select one representative edge per variant allele and remove redundant edges to limit the multiple testing burden. Edges are ranked by average depth across samples; the pipeline iteratively retains the top edge and discards those redundant with it (see Methods). Orange, retained representative edge; dashed gray, dropped edges. **(d)** Step 5: retain variable edges, defined as edges at which at least 10 samples carry a genotype differing from the majority. Red dashed boxes mark retained edges. **(e)** Step 6: for edges representing simple biallelic variants (e.g., >1>2), replace normalized edge depth with allele balance, calculated as the raw depth of the alternative allele edge divided by the summed depth of the alternative and reference edges. **(f)** Step 7: test edge quantification values (*i.e.*, allele balance or normalized edge depth) for association with traits.

We next performed a control experiment to evaluate EdgeDepth’s ability to measure genotype information using the HPRC2 pangenome graph. We applied EdgeDepth to short-read WGS data from the same HPRC samples used for genome assembly and graph construction, using a subset of samples (N=201) with long-read RNA-seq data from lymphoblastoid cell lines (LCLs) generated by the PacBio Kinnex platform^15^. Using the HPRC2 graph-based callset from the same 201 samples as the truth set, we evaluated the 1KG, PanGenie, and EdgeDepth callsets by whether each truth-set variant was represented in the callset (Methods). We found that EdgeDepth captures 97.4% of “common” variants (defined here as present in ≥10 samples) and 98.0% of lead eQTL variants that can be mapped using variant calls from the pangenome graph (**Fig. 2a**). This is a substantial increase over the published 1KG variant calls for these same samples (86.3% and 86.9%, respectively). As expected, the variant calling improvements are primarily in SVs and VNTRs, and in complex and repetitive parts of the genome (**Fig. 2b**). Notably, there are 115 eQTLs that are identified using the HPRC graph-based genotypes but not the 1KG dataset, and EdgeDepth captures 97 (84.3%) of these. In comparison, the most mature and widely used pangenome-based variant calling method, PanGenie^4^, identified 79% of variants, 77.9% of eQTLs, and 22.6% of graph-specific eQTLs, although we expect this to be a slight underestimate due to reference liftover effects, since PanGenie was run using the T2T-CHM13 reference^9^ and 2.8% of variants do not liftover to GRCh38 (see **Methods**). These results show that EdgeDepth is a sensitive and accurate method for measuring genotype information using pangenome graphs, that achieves superior trait association performance relative to the official 1KG callset and also the leading pangenome-based method.

**Figure 2.**
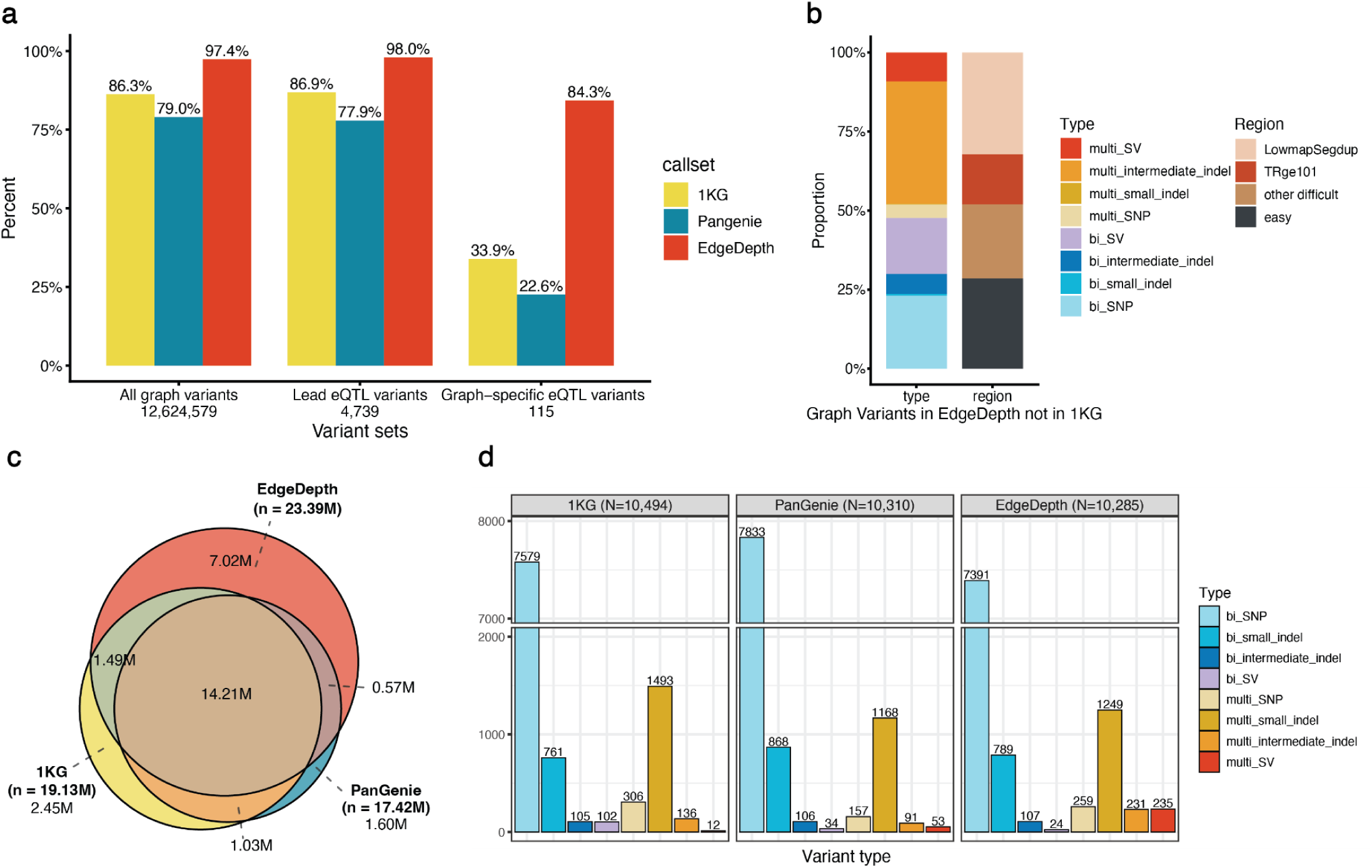
Comparison of variant calling and eQTL discovery. **(a)** Percentage of pangenome graph variants captured by each short-read WGS callset based on analysis of 201 HPRC samples. Bars are colored by callset and set sizes are shown below each group, including all variants in the graph, lead eQTL variants from graph-based eQTL analysis, and “graph-specific” lead eQTL variants that were not found by the 1KG analysis. **(b)** Composition by variant type (left) and genomic region (right) of the graph variants captured by EdgeDepth but not by 1KG (201 HPRC samples). Variant types are coded as bi_ or multi_ (biallelic/multiallelic) combined with SNP (1 bp), small indel (2-9 bp), intermediate sized indel (10-49 bp) or SV (≥50 bp). Genomic regions include low-mappability and segmental-duplication regions (LowmapSegdup), tandem repeats greater than or equal to 101 bp (TRge101), additional difficult regions not in those categories (other difficult), and easy regions. **(c)** Venn diagram of shared and unique variants across the 1KG, PanGenie and EdgeDepth callsets, based on 430 short-read WGS samples from AFGR. Variant counts are in millions, with per-callset totals in parentheses. **(d)** Distribution of variant types among lead eQTL markers for each callset, where N indicates the number of eGenes per callset. Note that the y-axis is broken between 2000-7000.

### Comparative eQTL mapping performance

We then applied EdgeDepth, PanGenie, and a third more specialized pangenomic method, danbing-tk^16^ to a separate set of 430 samples from 1KG that have deep (>30×) short-read WGS from 1KG^7^ and short-read RNA-seq data from LCLs generated by the African Functional Genomics Resource (AFGR)^17^. Most of these samples (407 of 430) are not included in the HPRC2 resource. PanGenie is a genome-wide method that can in principle genotype any type of variant represented in a pangenome graph, while danbing-tk is a specialized k-mer-based method designed specifically to genotype VNTRs. Both methods use k-mer counts rather than read alignment. After systematic filtering for variant quality and allele frequency, and retaining matched variants across callsets even when allele frequency measurements were slightly different (see **Methods**), the EdgeDepth callset includes 23.4 million variants, the PanGenie callset includes includes 17.4 million variants, the danbing-tk callset includes 22.0 thousand VNTRs, and the traditional reference-based 1KG callset includes 19.1 million variants (**Table 1, Supplementary Fig. 1**). There is strong overlap between the callsets in terms of shared variation, but also variants that are unique to one or two methods (**Fig. 2c**).

**Table 1.** Number of total variants and eQTL lead markers. Variant and eQTL counts are shown for the analyses where eQTL mapping was done separately for each variant callset (left) and jointly with pooled variants (right). Each row represents one variant callset. “No. variants” indicates the number of variants included in eQTL mapping, and “No. eQTLs” indicates the number of eQTLs identified, with the fraction of total shown in parentheses for the joint analysis.

| Variant callset | No. variants | No. eQTLs (separate) | No. eQTL lead markers (joint) |
| --- | --- | --- | --- |
| 1KG | 19,166,694 (31.9%) | 10,494 | 3,243 (31.7%) |
| PanGenie | 17,421,431 (29.0%) | 10,310 | 2,715 (26.5%) |
| EdgeDepth | 23,386,051 (39.0%) | 10,285 | 4,251 (41.6%) |
| danbing-tk | 21,962 (0.04%) | 541 | 14 (0.1%) |
| Total | 59,996,138 |  | 10,223 |

When applied separately each to their own eQTL mapping experiment, EdgeDepth and PanGenie show similar overall performance to the 1KG callset, identifying a similar number of genes with an eQTL (hereafter referred to as “eGenes”) overall (10,285 and 10,310, respectively, vs. 10,494) (**Table 1**). Within the 1KG callset, we observe that 75.1% of eQTLs are from SNPs, 23.8% from small indels, and 1.1% from SVs, which is roughly consistent with prior studies using similar methods^18–20^. The pangenome methods identify a roughly similar fraction of eQTLs with different variant types, except for a larger fraction of cases where the lead variant is a multiallelic intermediate sized indel (10-49 bp) or SV (≥50 bp) (**Fig. 2d**). For example, EdgeDepth identified 235 eQTLs (2.3% of total) where the lead variant is a multiallelic SV, whereas 1KG only identified 12 (0.1%). This indicates that the pangenome methods are providing high quality genotype information suitable for trait association, at a level that is roughly similar to the 1KG callset, and in addition are expanding the spectrum of putative functional variants that are identified as lead markers.

However, it is somewhat difficult to compare performance across separate eQTL mapping experiments because each variant callset has a different per-gene multiple testing burden (and hence p-value threshold using tensorQTL^21^), which benefits methods that detect fewer variants, and because most eQTLs can be discovered from linked proxies via linkage disequilibrium (LD) even when the most strongly associated and potentially causal variant is not present in the callset.

We thus performed a joint eQTL mapping experiment that included all variants identified by all 4 methods, allowing for multiple genotype representations per variant. This approach allows the methods to compete with each other to identify the most significant association for each gene, using a single per-gene significance threshold. A given method could identify the lead variant either by having identified a unique variant not found by the other methods, or by having the most accurate genotype information for its version of the lead variant. EdgeDepth identifies the most lead markers in this experiment (41.6%), followed by 1KG (31.7%) and PanGenie (26.5%) (**Table 1**). Whereas the vast majority of eQTLs have significantly associated variants from 2 or more methods (**Fig. 3a**), in most cases including 1KG and at least one pangenome-based method, there are 222 eQTLs (2.2% of total) that were identified solely by pangenome-based methods. Perhaps more importantly, given that exceeding the eQTL p-value significance threshold is arbitrary when considering power improvements that generalize to GWAS, there were 185 eQTLs (1.5% of total) for which the pangenome-based methods had a “strong” power boost (50% improvement in p-value) over the 1KG callset, and 812 (7.9%) that had a “moderate” boost (20% improvement). EdgeDepth was by far the most valuable method for improving association significance at complex loci: of the 185 eQTLs that showed a strong power boost, 85.9% were detected at the more significant level by EdgeDepth, 20.0% by PanGenie, and 4.3% by danbing-tk (**Fig. 3b**).

**Figure 3.**
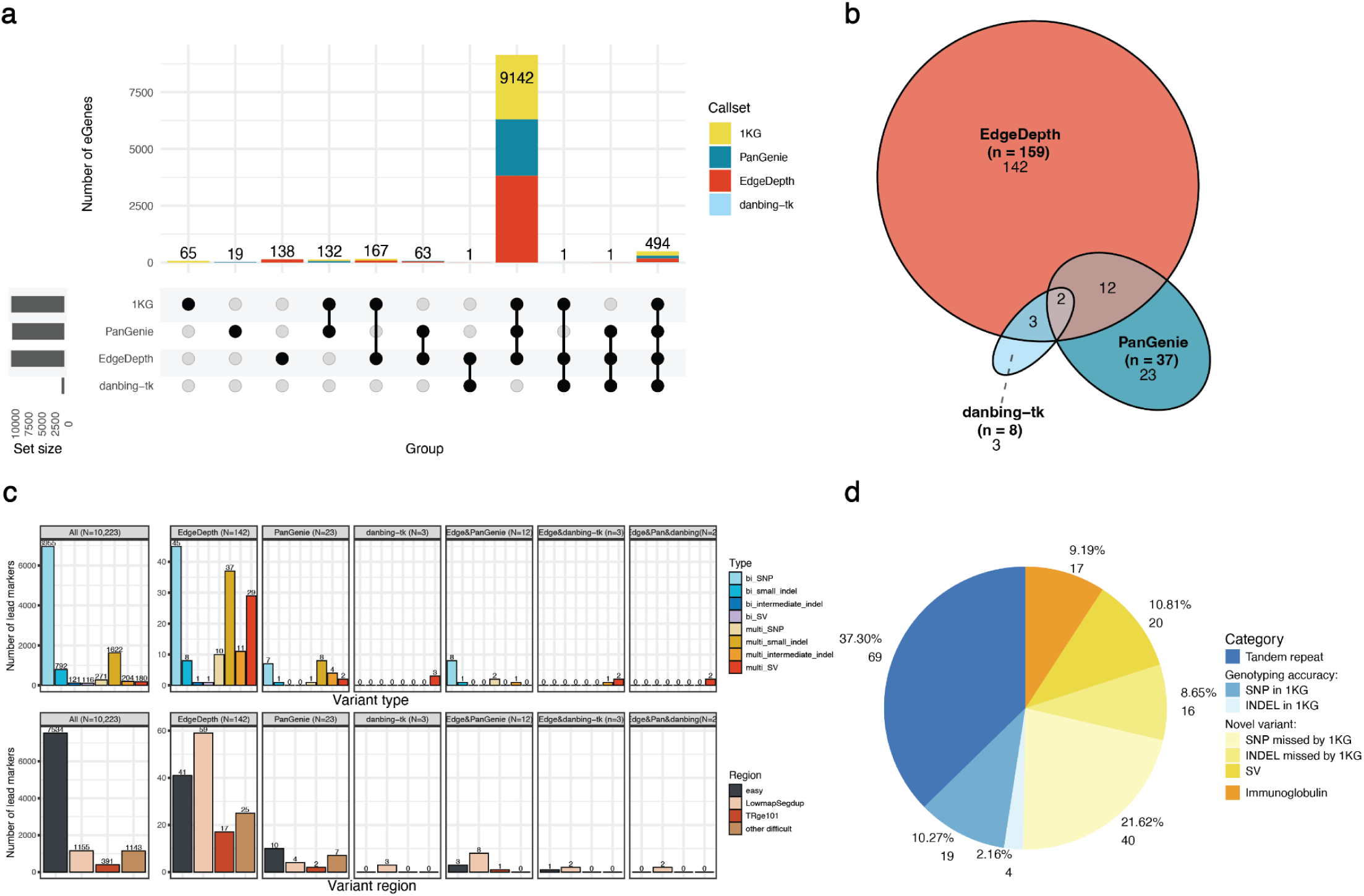
Joint eQTL analysis. We mapped eQTLs in the 430 AFGR samples using variants pooled from the 4 callsets, and characterized the lead markers at which pangenome variants strongly improved the eQTL signal (i.e., showed a strong “power boost”). **(a)** UpSet plot of eGenes identified by callsets. A callset detects an eGene if it contributed at least one variant with a nominal p-value below the gene-specific threshold determined by tensorQTL. eGenes are colored by lead marker source callsets. Panels b-d are focused on the 185 eGenes at which pangenome variants strongly improved the eQTL signal, defined as a >50% increase in -log10(nominal p-value) for the most significant pangenome variant relative to the most significant 1KG variant. **(b)** Euler diagram assigning the 185 eGenes to the pangenome callsets (PanGenie, EdgeDepth, and/or danbing-tk) that drove the improvement, defined by if the callset reported any variant with >50% of improvement. **(c)** Composition of lead eQTL markers by variant type (top) and genomic region (bottom). The left panel shows all eGenes from the joint analysis (N=10,223). The right panels show the 185 improved eGenes split by the driving callset group as in (b). Variant type abbreviations are as in Fig. 2. **(d)** Pie chart that breaks the 185 improved eGenes into categories that might explain the reason for improved p-value, including (from top) whether the lead variant is: a tandem repeat; a SNP or indel that was detected by 1KG; a SNP or indel that was missed by 1KG; an SV; or present in a complex immunoglobulin region.

Inspection of the 185 eQTLs with a strong power boost revealed that most are multiallelic SVs and indels, many of which are found in tandem repeats, segmental duplications, and other difficult to analyze genomic regions (**Fig. 3c**). For example, 20.5% are multiallelic SVs and 33.5% are multiallelic indels, whereas these variants represent merely 1.8% and 17.9% of total eQTLs, respectively. These same patterns are found in the 812 eQTLs with a more moderate power boost, albeit with somewhat weaker enrichment of multiallelic SVs (**Supplementary Fig. 2**). Of the strongly boosted eQTLs, 70.3% are found in difficult genomic regions comprised of segmental duplications, tandem repeats, and other repetitive sequences, as compared to 26.3% in the overall dataset. We classified the mechanisms underlying 185 strongly boosted eQTLs into one of four mutually exclusive categories: tandem repeats, which the methods employed by 1KG (i.e. GATK plus various SV detection tools) handle less effectively (37.3%); novel variants found in one or more pangenome callsets but not 1KG (41.1%); differences in genotyping accuracy (12.4%) and immunoglobulin (IG) genes (9.2%). IG genes were classified separately because this study used LCLs, in which IG loci should be interpreted cautiously due to somatic rearrangement **(Fig. 3d)**.

Among these loci there are many noteworthy examples that demonstrate the real-world value of using pangenomic methods, including 34 medically relevant genes^22^. For example, at the *MINPP1* gene, an 122 kb inversion identified by EdgeDepth is strongly associated with increased expression, apparently by rearranging the order of enhancers 75 kb upstream of the gene promoter (**Supplementary Fig. 3a**). This inversion is not identified in the 1KG or PanGenie callsets, likely because the breakpoints lie within segmental duplications in the reference, and the most strongly associated linked marker in 1KG is ∼20 orders of magnitude less significant (p=7.87×10^-41^ vs. 4.74×10^-21^). At the *SURF1* gene, EdgeDepth identified an intronic VNTR associated with increased expression, where different haplotypes carry 1-3 extra copies of a 7 bp repeat (**Supplementary Fig. 3b**). Interestingly, although the expansion alleles are identified by 1KG and PanGenie, they are represented as three distinct biallelic variants, whereas one of the edges measured by EdgeDepth captures the difference between zero versus one or more extra copies, and results in a vastly more significant p-value (p=2.41×10^-84^ vs. 5.29×10^-26^). This example demonstrate a key strength of the graph-based association approach, where graph features can represent structural relationships among complex variant alleles in ways that are difficult to capture from testing allele dosage. A similar example is found at the *TRIB3* gene, where EdgeDepth and danbing-tk (but not 1KG) identified a promoter VNTR with a 33 bp repeat unit present in 1 to 10 copies, at which expansion is linked to increased expression (**Supplementary Fig. 4a**). Notably, the regulatory effects of this VNTR on *TRIB3* have been independently validated by luciferase reporter assays^23^. A more simple example is found at the *CBS* gene, where EdgeDepth and PanGenie identified a SNP in exon 10 that is not reported in the 1KG callset, and that is much more strongly associated with gene expression than the linked SNP reported by 1KG (p=3.22×10⁻^21^ vs. 8.93×10⁻^9^, **Supplementary Fig. 4b**). 1KG appears to have missed this variant due to a false segmental duplication in GRCh38 that creates *CBSL*, an incorrect copy of *CBS*, causing reads to misalign to *CBSL* instead of *CBS*, and resulting in false negative variant calls across the *CBS* locus^22^.

However, it is also important to note that the 1KG callset outperforms the pangenome callsets at many eQTLs (**Supplementary Fig. 5**), usually by a small margin but sometimes more substantially (*e.g.*, 108 eQTLs are >50% better in 1KG), indicating that current pangenomic methods have ample room for improvement.

Moreover, although EdgeDepth found the most lead variants in the joint analysis, this is partially due to the fact that PanGenie and 1KG often have identical genotype information, and ties are split between them, whereas EdgeDepth represents variants in a different way. Indeed, in head-to-head comparison at shared variants detected by multiple methods (in effect measuring genotype quality), both 1KG and PanGenie slightly outperform EdgeDepth in terms of the number of eQTL lead variants identified, with PanGenie performing best (**Supplementary Fig. 6**). Taken together, the results indicate that PanGenie and 1KG typically provide higher quality genotype information at the variants that they are able to detect, likely due to the fact that they represent easier-to-detect biallelic variants using discrete genotypes rather than noisier allele balance values, whereas EdgeDepth performs less well at simple variants but enables the assessment of novel complex variation via thorough interrogation of graph features. We further note that there are additional variant calling methods currently under development that, to varying degrees, use pangenomic approaches (e.g., DeepVariant^24^, DRAGEN^25^, and PanVariants^26^), that were not tested here due to practical constraints related to 1KG callset availability and pipeline maturity relative to the HPRC2 release. Further evaluation of best-in-class pangenomic methods using HPRC2 and future pangenome releases will be an important area of future work.

### Added benefit of pangenome-specific variation to eQTL discovery and fine-mapping

A realistic near-term use-case for pangenomic tools may be to supplement traditional workflows by adding novel pangenome-specific variants. This approach is designed to leverage the best features of different available methods, in effect combining the high-fidelity genotypes provided by current reference-based tools (e.g., GATK) at easier-to-detect variants with the complementary ability of pangenomic tools to ascertain additional more complex variant types. To test this, we created a merged variant callset based on the nonredundant union of all four variant callsets used in our study, retaining the 1KG version in cases of redundancy, and compared the performance of the merged “HPRC2+1KG” callset to 1KG (**Table 2**). We performed eQTL mapping with the merged callset and found 10,169 eQTLs, which is roughly similar to the number found using the 1KG callset (10,494). However, the spectrum of putative functional variants is quite different in the merged callset, with 20.3% of lead variants contributed by the pangenome-based methods, and (similar to described above) a much larger fraction of multiallelic SVs and indels (**Fig. 4a**). Moreover, the merged callset significantly improves power at a significant number of eQTLs, with 893 eQTLs (8.8% of total) showing a ≥20% improvement in p-value and 581 (5.7%) showing a >50% improvement relative to the 1KG callset alone (as described in the companion HPRC2 paper^1^. These results are consistent with and complementary to those obtained via the separate and joint eQTL mapping experiments, and further support the added value of ascertaining complex forms of genetic variation for trait association.

**Figure 4.**
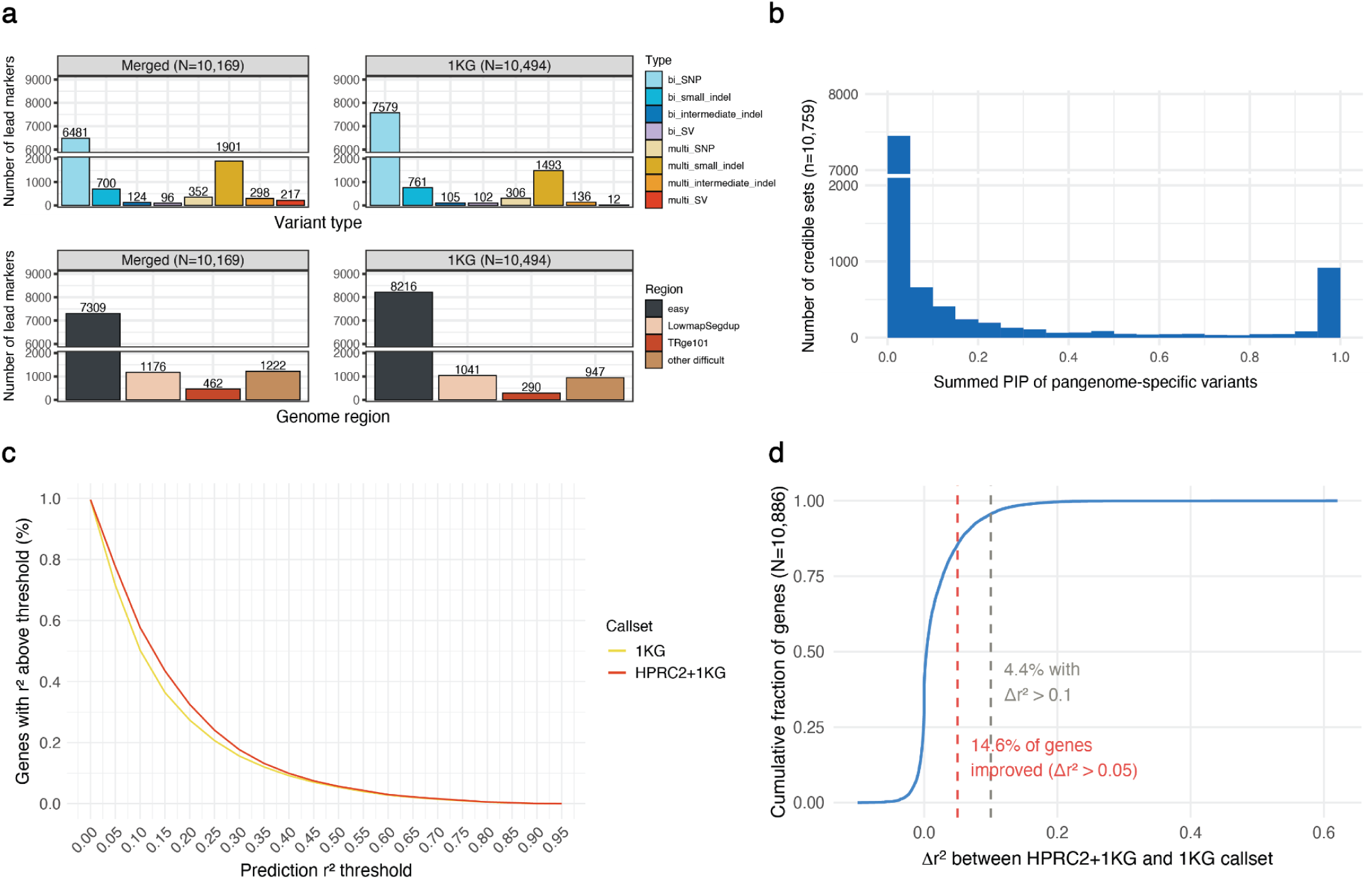
Analysis of the merged HPRC2+1KG callset. We compared lead eQTL marker composition, fine-mapping signals, and gene expression prediction accuracy between the 1KG callset and a merged callset composed of 1KG variants plus pangenome-specific variants. **(a)** Composition of lead eQTL markers by variant type (top) and genomic region (bottom) for the merged callset (left) and 1KG alone (right). N indicates the number of lead markers per callset. Variant type abbreviations are as in Fig. 2. **(b)** Distribution of the summed per signal posterior inclusion probability (PIP) of pangenome variants in each credible set (CS, N=10,759), from fine-mapping the merged callset with SuSiE^27^. **(c)** Percentage of 10,886 union eGenes whose expression is predictable above a given r^2^ threshold (x axis) for the merged HPRC2+1KG and 1KG callsets. Lines are colored by callset. For each gene and callset, expression was predicted using variants within 1 Mb of the transcription start site, and quantified by the 5 fold cross-validated r^2^ between predicted and observed expression, taking the highest adjusted r² across five methods (Lasso, ridge, elastic net, top-1 single best variant, and SuSiE). **(d)** Cumulative distribution of Δr^2^ (r^2^ of merged HPRC2+1KG callset - r^2^ of 1KG) across 10,886 genes. Positive values indicate improved expression prediction with the merged callset. 14.6% of genes improve by Δr^2^ > 0.05 (red dashed line) and 4.4% by Δr^2^ > 0.1 (gray dashed line).

**Table 2.**
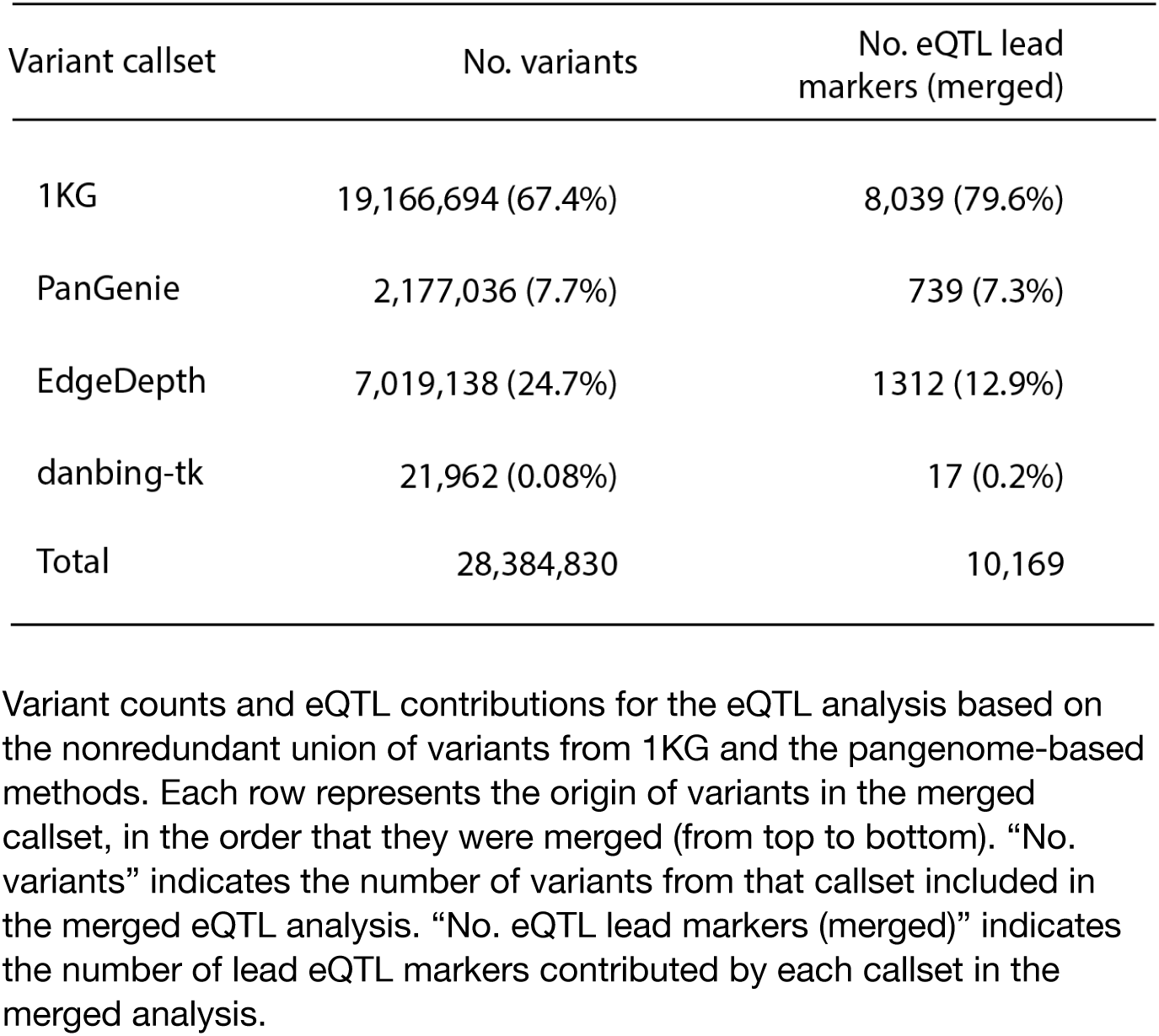
eQTL results using the merged HPRC2+1KG callset. Variant counts and eQTL contributions for the eQTL analysis based on the nonredundant union of variants from 1KG and the pangenome-based methods. Each row represents the origin of variants in the merged callset, in the order that they were merged (from top to bottom). “No. variants” indicates the number of variants from that callset included in the merged eQTL analysis. “No. eQTL lead markers (merged)” indicates the number of lead eQTL markers contributed by each callset in the merged analysis.

| Variant callset | No. variants | No. eQTL lead markers (merged) |
| --- | --- | --- |
| 1KG | 19,166,694 (67.4%) | 8,039 (79.6%) |
| PanGenie | 2,177,036 (7.7%) | 739 (7.3%) |
| EdgeDepth | 7,019,138 (24.7%) | 1312 (12.9%) |
| danbing-tk | 21,962 (0.08%) | 17 (0.2%) |
| Total | 28,384,830 | 10,169 |

To explore these association signals in more detail, we performed fine-mapping to predict causal variants using SuSiE^27^. In contrast to standard eQTL mapping, which reports the single most strongly associated “lead” variant for each eGene, fine-mapping allows for multiple association signals per gene, and provides probabilistic causal variant prediction for each signal by integrating association signal strength and LD structure. Using the merged HPRC2+1KG callset, we identified 10,759 credible sets (CSs) in 7,171 eGenes (**Supplementary Fig. 7**), where a CS is defined to encompass 95% of the posterior inclusion probability (PIP) for a given association signal. Among these, 40.0% of CSs and 49.2% of genes included at least one pangenome-specific variant, and pangenome-specific variants account for 19.3% of the 2,965 high-confidence causal variants with posterior probability α > 0.95. These results are consistent with the eQTL results, and suggest that pangenome-specific variants do not merely add additional correlated signals but contribute to the nomination of putative causal variants for a substantial fraction of genes.

We next compared the fine-mapping results from the HPRC2+1KG callset to the 1KG callset alone. Among 6,893 eGenes with one or more CSs identified in both analyses, we classified signals based on how PIP shifted after adding pangenome-specific variants. At 937 (13.6%) genes, pangenome variants account for the majority of summed PIP (>0.5) in at least one CS, indicating pangenome variants refined the signal and help prioritize a candidate causal variant (**Fig. 4b**). These include 315 association signals that were also found by 1KG, and 856 novel signals that were only found by pangenome-specific variants, where novel signals notably represent 8.0% of total CSs (see **Supplementary Fig. 8** for examples). Of the 937 genes, there were 166 where a high-confidence causal variant (α>0.95) was nominated by the merged callset but not by the 1KG callset, and 253 where 1KG had nominated a high-confidence causal variant that turned out to be incorrect. In contrast, at 15.9% genes, pangenome variants contributed a more moderate (0.1-0.5) level of PIP to one or more CSs, and at 4,692 (68.1%) genes, pangenome variants did not contribute meaningfully and the association signals remained stable. At 3.7% of genes, including pangenome variants resulted in lost CSs, apparently by increasing the number of variants in LD and causing a loss of “purity” in CS composition.

### Gene expression prediction

Another measure of utility is whether a variant callset improves the ability of predictive models to explain gene expression variation in unseen samples. Transcriptome-wide association study (TWAS) methods such as FUSION^28^ use multiple types of linear models to predict gene expression based on variants in a cis window (e.g., 1 Mb) around the gene and have proven to outperform best-in-class deep learning models^29,30^. We trained LASSO, ridge regression, elastic net, top-eQTL models implemented in FUSION, and SuSiE, to perform TWAS-style gene expression prediction on the union of eGenes (N=10,886) found using the 1KG and HPRC2+1KG callsets. The merged HPRC2+1KG callset exhibits improved prediction across the full set of genes (**Fig. 4c, Supplementary Fig. 9**), with a higher mean (17.6% vs. 15.8%) and median (12.5% vs. 10.1%) variance explained relative to 1KG. These include strong improvements at a subset of genes, including 1,588 genes (14.6%) with an improved r^2^ of at least 0.05, and 474 genes (4.4%) with an improved r^2^>0.1 (**Fig. 4d**). This result is important because it shows that the additional variants added by pangenome-based variant detection methods in the HPRC2+1KG callset – often complex multiallelic variants in repetitive regions – are functionally relevant and adding useful information for predictive modeling across a large fraction of genes. These benefits will be increasingly important as efforts to employ deep learning models for predicting haplotype-level functional effects continue to improve.

### Colocalization with GWAS loci

Finally, we explored the disease relevance of the eQTLs reported here. We examined the 812 eQTLs with a strong or moderate pangenome power boost to see if any of these associations implicated a novel causal variant that could explain prior GWAS results. We performed colocalization analysis using the *coloc* software^31^ and a set of 41 studies from the GWAS catalog^32^ (**Supplementary File 1**) with harmonized summary statistics from diverse traits including blood biomarkers (e.g. lymphocyte count), common complex disease (e.g. Type 2 diabetes), and conditions related to immune function (e.g. Crohn’s disease) (as in AFGR study^17^). We ran *coloc* on 156 gene-variant pairs found within 100 kb of a GWAS lead marker, and identified 10 cases where both the eQTL and the GWAS trait share a single causal variant (defined as H4≥0.5) (**Fig. 5, Supplementary Figs. 10-14, Supplementary File 2**). We note that this is likely to be an underestimate of the yield expected from similar future studies given that our eQTL mapping population is 100% African ancestry, whereas the GWAS populations are primarily European ancestry. All of these examples involve multiallelic variants or difficult-to-analyze genomic regions. Two examples are at the *HLA-A* locus, where a single multiallelic SNP in an HLA-A exon affects expression of *HLA-A* and the *HCG4P5* pseudogene, and is associated with multiple sclerosis. Three examples involve intronic short tandem repeat (STR) length variants, including effects on *UBE2R2* and Type2 diabetes, *CREM* and Crohn’s disease, and *PPM1G* and triglyceride levels.

**Figure 5.**
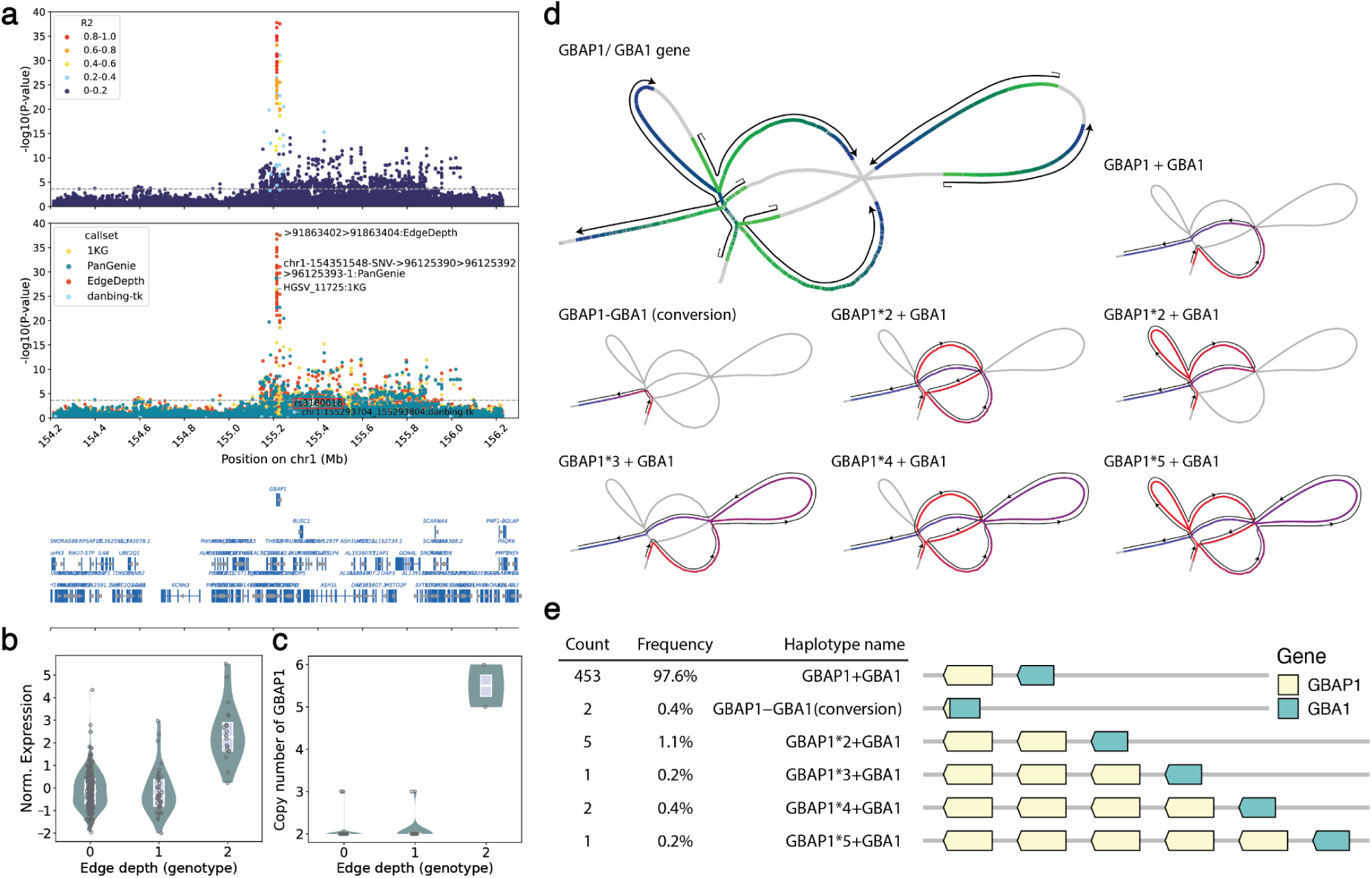
The *GBAP1* eQTL that colocalizes with a GWAS locus for Chron’s disease. **(a)** LocusZoom plot of the *GBAP1* locus within 1 Mb of the transcription start site (TSS); y-axis, −log_10_(nominal p-value); x-axis, GRCh38 position on chr1. Top, variants colored by r^2^ (LD) to the lead eQTL marker; middle, variants colored by callset, with the lead marker of each callset annotated and the previously reported lead variant (rs3180018) marked by a red rectangle; bottom, gene annotations in the window. The dashed line marks the gene-specific nominal p-value significance threshold. **(b)** Normalized *GBAP1* expression versus the EdgeDepth lead-marker copy number across 430 AFGR samples. Edge depth genotype is called from fitting a constrained Gaussian mixture model (GMM) from genomeSTRiP^75^ to continuous edge depth dosage. Boxes show the median and interquartile range (25th-75th percentile). Individual samples are shown as points. **(c)** Diploid *GBAP1* copy number versus lead-marker genotype in the 201 fully assembled HPRC samples. **(d)** Structural haplotypes of the *GBA1*/*GBAP1* locus across 464 HPRC2 assemblies. Top left, gene-copy locations (color gradient green to blue marks the start and end of each copy); remaining panels, the graph path of each structural allele (gradient red to blue from path start to end). **(e)** Linear visualization and frequency of the structural haplotypes in d.

Perhaps most interestingly, our results implicate *GBAP1* pseudogene copy number in Crohn’s disease through a novel mechanism that to our knowledge has not been previously reported for a common disease GWAS locus (**Fig. 5**). Prior studies identified a Crohn’s disease GWAS peak near the *GBA1* gene (the original candidate genes were *SCAMP3* and *MUC1*)^33^ and showed via fine-mapping and colocalization analyses that this association is due to the same causal variant that underlies an eQTL for *GBAP1* in four T-cell types^34^, where the disease risk variant increases *GBAP1* expression. Here, we identify this same *GBAP1* eQTL and also observe colocalization (**Supplementary Fig. 15**), but in our case EdgeDepth identifies pangenome-specific paralogous sequence variants (PSVs) within the *GBAP1* gene body that are >20 orders of magnitude more strongly associated with *GBAP1* expression than the lead eQTL variant identified previously (rs3180018) (**Fig. 5a,b**). Fine-mapping with SuSiE prioritized a pangenome edge variant “>91863422>91863424” as the most likely causal variant, with a posterior inclusion probability (PIP) of 0.9998, and a 95% credible set containing a single variant. Alignment depth at these PSVs reflects the dosage of a specific *GABP1* version that is not present in the GRCh38 reference genome (**Fig. 5c**), and based on the 462 HPRC assemblies, individual haplotypes carry 1-5 copies of *GBAP1* pseudogene sequences at this region (**Fig. 5d,e**). Remarkably, there is experimental evidence to indicate that *GBAP1* transcripts function as a competing endogenous RNA (ceRNA) “sponge” for the miR-22-3p microRNA that downregulates regulates *GBA1*^35^, providing a plausible mechanism by which increased *GBAP1* pseudogene copy number leads to increased expression of the *GBA1* gene. Although more work is required to unequivocally demonstrate that this specific molecular mechanism explains the Crohn’s Disease association at this locus, our data provide evidence that it does, and further point to *GBA1* transcript levels as the underlying cause of disease risk and a logical target for drug discovery efforts.

## Discussion

Taken together, the results presented here demonstrate that, relative to traditional reference-based methods, newly-developed pangenome-based variant calling methods enable more powerful analyses of genetically-regulated gene expression variation by providing meaningful improvements to trait association, fine-mapping, and prediction. As we show above using *GBAP1* and other examples, these quantitative improvements translate into novel genetic findings, such as the identification of new candidate causal variants and the discovery of new potential disease mechanisms based on colocalization of eQTL and GWAS signals.

Our development of the EdgeDepth method represents a novel strategy for pangenome-based trait association. Rather than attempting to resolve allelic structure and assign discrete genotype combinations to each sample in a study – which remains intractable at many complex loci – EdgeDepth bypasses this requirement and performs trait association directly on the graph itself. This allows arbitrarily complex alleles to be tested efficiently, requiring only that one or more informative markers for each allele has been captured in the pangenome graph, and that WGS data can be aligned to these markers in a relatively accurate way. As pangenome resources continue to grow, and graph-based read alignment continues to improve, EdgeDepth and related methods will become even more powerful. A weakness of EdgeDepth is that the lack of hard genotype calls and copy number dosage values limits the utility for population genetics and related areas, and may slightly diminish trait association power at easy-to-genotype variants due to increased noise.

Beyond gene expression variation, we expect these findings to extend to future pangenome-based association studies of common diseases and other complex traits. Although there are important differences between bulk-tissue eQTLs and GWAS loci in terms of cell-type specificity, effect sizes, and the specific association signals that are detected^36–39^, the vast majority of GWAS hits are nonetheless caused by common genetic variants that alter gene expression, and it is difficult to imagine that the spectrum of trait-associated variation would be markedly different in terms of variant types and the ability of different methods to detect and genotype them accurately. Our results therefore suggest that application of pangenome-based methods to GWAS will provide broadly similar improvements to trait association and fine-mapping, and hence lead to the discovery of new variants and genes underlying complex traits. A key challenge moving forward will be to improve the cost and scalability of graph-based genome analysis, so that these methods can be affordably applied at the scale of modern studies.

## Methods

### The EdgeDepth pipeline

#### Alignment and normalization

We aligned short-read WGS data from the 1KG to the HPRC2 Minigraph-Cactus pangenome graph using `vg giraffe` (v1.64.1)^13^ with haplotype-sampling. The graph file (.gbz) and haplotype information (.hapl) were provided by the HPRC2. We generated k-mer counts of each sequencing sample using the kmc tool (v3.2.4)^40^ (`kmc -k29 -koff`) then performed haplotype sampling (integrated in Giraffe) and alignment using `vg giraffe` with default parameters. For each sample, we quantified read support for each graph edge from the resulting alignments using `vg pack`. To account for differences in sequencing depth and edge-depth composition across samples, we normalized edge depths using DESeq2’s median-of-ratios method^41^. For each edge, we calculated the geometric mean depth across samples to define a pseudo-reference profile. For each sample, we then calculated the median value of all ratios between the sample’s edge depths and the corresponding pseudo-reference values, and took it as a sample-specific size factor. Edges with zero depth in one or more samples were excluded from size-factor estimation. Normalized edge depth was calculated by dividing each sample’s raw edge depth by the corresponding size factor.

#### Selection of representative edges

A given variant allele can be represented by multiple edges in the pangenome graph. To reduce redundancy and multiple-testing burden, we selected a single representative graph edge with the greatest read-depth support per variant allele. To improve computational efficiency, we first decomposed the pangenome graph into biconnected component subgraphs. This allowed redundant-edge filtering to be performed within smaller locally connected variant structures rather than across the full graph. Within each subgraph, we considered only non-reference edges and ranked them by average raw depth across all samples. Proceeding from the highest to lowest depth edge, we iteratively selected each edge as a representative, removed it from the graph, and identified its redundant edges as follows: upon removing the representative edge from the subgraph, any newly formed bridge edges were designated as its redundant edges and removed. This criterion reflects the property that every cycle containing the representative edge also contains its redundant edges, ensuring that the representative edge uniquely tags the corresponding allele. This procedure was repeated until no non-reference edges remained in the subgraph. For large subgraphs where exhaustive search was computationally expensive, we applied a local approximation that searched for redundant bridge edges within 50 graph steps of the selected edge. The final set of representative edges were used as nonredundant allele markers for downstream analyses.

#### Alignment depth filters for candidate variable edges

We applied read-depth filters to retain edges with sufficient read support and remove unreliable edges that arise from highly repetitive regions. We retained edges with a raw depth of at least 5 reads in at least 10 samples. We also removed edges with an average raw depth of at least 1,000 across samples, because extremely high-depth edges are likely to occur in centromeric or other highly repetitive regions where short-read alignment is less reliable.

#### Selection of variable edges

To identify and retain edges that show variation across the short-read WGS samples, we fitted a constrained Gaussian mixture model (GMM) to the raw read depth distribution of each representative edge passing alignment depth filters. The model followed the Genome STRiP^42^ depth clustering framework used for CNV detection, in which genotype clusters are modeled by Gaussian components with linearly constrained means and variances. The number of mixture components was set to the maximum observed depth divided by the estimated single copy mean depth (rounded up to the nearest integer). An edge was classified as variable if the fitted GMM supported a non-major genotype group containing at least 10 samples. Edges with insufficient depth to fit more than one component were also removed.

To recover edges incorrectly rejected by the GMM filter, we applied a rescue step for edges that failed the above criterion but showed at least 10 outlier depth values. This step was motivated by cases in which a near-zero depth peak was slightly shifted above zero, causing the GMM to fit the near-zero cluster as the primary component rather than a true nonzero genotype group. An edge was rescued if at least 10 samples had depth exceeding median ±4 median absolute deviations (MAD), and if the upper outlier threshold (median + 4 MAD) exceeded an estimated heterozygous depth cutoff (4.71). The heterozygous depth threshold was estimated by fitting the constrained GMM to a random 0.001% subsample of all edges after excluding extreme high-depth tails (>68) and the dominant zero-depth edges. From the distribution, we identified the heterozygous peak with mean 16.99 and standard deviation 6.14, and defined the heterozygous depth cutoff as the mean minus two standard deviations.

#### Allele balance calculation

To more accurately represent genotype dosage at biallelic variants, we replaced normalized edge depth with allele balance (AB) values based on read depth information from multiple edges. A variant was defined as biallelic based on graph topology: specifically, a top-level snarl anchored at both ends to the reference path, containing exactly two traversals and no nested snarls. This biallelic variant definition encompasses SNPs, simple insertions, and simple deletions regardless of variant lengths. For each biallelic variant, AB was calculated as raw depth of the alternative allele edge divided by the sum of the alternative and reference allele edge depths. Among the two reference edges representing the reference allele, we selected the reference edge with higher average raw depth for AB calculation. The resulting AB value represents the relative dosage of the alternative allele across all samples, ranging from 0 (homozygous reference) to 1 (homozygous alternative). For edges passing all filters, AB was used as the genotype dosage for biallelic variants, while normalized depth was used for all remaining edges. These values were used as variant dosages in downstream eQTL analyses. The complete EdgeDepth pipeline, from input sequencing reads to final variant table generation, is available at https://github.com/lushjia/EdgeDepth.

### Annotation of graph edges

#### Assignment of GRCh38 coordinates to graph edges

To represent edges in VCF format with linear reference coordinates for cis-eQTL analysis, we assigned each graph edge a GRCh38 coordinate. Because an edge is defined by two connected graph nodes, we first assigned GRCh38 coordinates to individual nodes. Nodes in the reference path were assigned coordinates from the SO tag annotations in the graph rGFA file. For non-reference nodes, we traversed upstream and downstream neighbors in the graph using libbdsg (v0.4)^43^ until reaching reference nodes. Because a non-reference node can connect to multiple upstream or downstream reference nodes, we summarized its reference coordinates by the last base of the leftmost upstream reference node and the first base of the rightmost downstream reference node. Nodes without an identifiable upstream or downstream reference neighbor were assigned missing coordinates on the corresponding side.

Edge coordinates were then assigned based on node coordinates. For each edge, if the node with a smaller identifier (indicating a more upstream genomic position) was on the GRCh38 reference path, the edge coordinate was defined as the last base of that node. If the node with a smaller identifier was not on the reference path, we used its upstream reference coordinate. When this coordinate was missing, we used its downstream reference coordinate. Edges for which both upstream and downstream reference coordinates were missing were processed with the `get_vcf_position` function from the Pantree package (v0.1.0)^44^ to recover a coordinate. The resulting edge reference coordinate was used to construct a VCF representation of the edge depth matrix for eQTL analysis.

#### Variant allele annotation

Edges in the pangenome graph represent variants, but they are not automatically interpretable as variant alleles or sequences. To enable biological interpretation of edge based measurements, we mapped each edge to the variant alleles it could represent based on chromosomal position, alternative alleles and corresponding reference alleles.

Variant alleles were identified by tracing HPRC haplotype paths through the graph and recording deviations from the reference path. Specifically, for each edge, we identified all alternative traversals passing through that edge. We defined an alternative traversal as a haplotype path segment that deviated from the reference path and rejoined it at downstream reference sequence. We located the nearest upstream and downstream reference nodes flanking the deviation. The alternative allele sequence was extracted from the alternative traversal between these two anchor nodes, and the reference allele sequence was extracted from the corresponding reference path between the same anchors. Chromosomal coordinates were assigned based on the GRCh38 coordinate of the upstream reference anchor node. Alleles were normalized and left-aligned using `bcftools norm` (v1.21)^45^ with GRCh38 FASTA (`-f -c x`). The resulting annotation links each edge to all variant alleles represented by alternative traversals containing that edge. A single edge can map to multiple alleles and a single allele can involve multiple representative edges.

### Genomic datasets and variant callsets

#### Cohorts

This study used two cohorts. The first cohort comprised 201 samples selected from 206 HPRC samples with PacBio Kinnex long-read RNA sequencing data^1^. Five samples were excluded: one (HG00272) because it was not included in the HPRC2 Minigraph-Cactus graph construction, and four additional samples (HG01123, HG02486, HG02559, HG03471) because they were absent from the 3,202 1KG samples with high-coverage short-read WGS data^7^. This cohort was used to evaluate the ability of multiple short-read based variant callsets to capture graph based variant calls. The second cohort comprised 430 samples of five African populations, including 99 Esan in Nigeria (ESN), 112 Gambian in Western Division, The Gambia - Mandinka (GWD), 97 Luhya in Webuye, Kenya (LWK), 83 Mende in Sierra Leone (MSL), 39 Yoruba in Ibadan, Nigeria (YRI), derived from the overlap between 593 samples from the African Functional Genomics Resource (AFGR)^17^ and the 3,202 1KG samples. Of the 433 overlapping samples, three were excluded: two (NA18489 and NA18874) were absent from the quality control results provided by 1KG on the Terra platform, and one (NA18487) was absent from the DNA sequencing data (European Nucleotide Archive accession PRJEB31736). This cohort was used for eQTL discovery and comparison across short-read based variant callsets.

#### Short-read WGS-based variant callsets

We curated 4 variant callsets and evaluated their performance in eQTL discovery based on the number of eQTLs identified and signal strength. The callsets include one GRCh38-based callset from 1KG and three pangenome reference-based callsets: PanGenie, danbing-tk, and EdgeDepth (described in the EdgeDepth pipeline section).

#### 1KG callset

We downloaded 1KG variant calls including small variants genotyped by GATK (https://ftp.1000genomes.ebi.ac.uk/vol1/ftp/data_collections/1000G_2504_high_coverage/working/20201028_3202_phased) and structural variants integrated from multiple SV callers^7^ (downloaded from http://ftp.1000genomes.ebi.ac.uk/vol1/ftp/data_collections/1000G_2504_high_coverage/working/20210124.SV_Illumina_Integration/1KGP_3202.gatksv_svtools_novelins.freeze_V3.wAF.vcf.gz). We extracted genotypes for the 201 HPRC and 430 AFGR samples separately and excluded variants with minor allele sample (MAS) count less than 10 in each cohort.

#### PanGenie callset

Variants of 3,202 1KG samples were genotyped with decomposed vcf derived from the HPRC2 T2T-CHM13-based Minigraph-Cactus graph using PanGenie (v4.2.1)^4^ (https://s3-us-west-2.amazonaws.com/human-pangenomics/pangenomes/scratch/2026_03_30_pangenie/pangenie_all-samples_filtered.vcf.gz). The PanGenie callset covers both small variants and SVs. We ran `picard LiftoverVcf` (v2.25.6, https://broadinstitute.github.io/picard/) to lift over variant coordinates from T2T-CHM13 to GRCh38. The T2T-CHM13 to GRCh38 chain was downloaded from the UCSC genome browser (https://hgdownload.gi.ucsc.edu/hubs/GCA/009/914/755/GCA_009914755.4/liftOver/chm13v2-hg38.over.chain.gz). We set `--RECOVER_SWAPPED_REF_ALT` true to rescue variants where the T2T-CHM13 ALT allele matched the GRCh38 REF allele, which swaps the REF and ALT alleles accordingly. Of the 40,167,722 total variants, 1,144,369 (2.8%) failed liftover and were excluded. We extracted genotypes of 201 HPRC and 430 AFGR samples separately and removed variants with MAS count below 10 in each cohort.

#### EdgeDepth callset

We applied EdgeDepth pipeline to call genetic variants, including small variants and SVs, for 201 HPRC and 430 AFGR samples using the HPRC2 GRCh38-based Minigraph-Cactus graph^1^. Steps included alignment to the pangenome graph, depth normalization, edge filtering, and allele balance estimation, as described in the EdgeDepth pipeline section. Variants were filtered in each cohort as described above.

#### danbing-tk callset

This callset quantified tandem repeat dosage using danbing-tk^16^ along with its bias correction submodule (commit `4b55315`). We extracted tandem repeats of 430 AFGR samples and excluded tandem repeats that lacked GRCh38 coordinates, had non-missing genotypes in fewer than 10 samples, or had a dosage of zero across all 430 samples.

#### Pangenome graph-based variant callset for the 201 HPRC samples

All variants of HPRC2 input samples are embedded in HPRC2 GRCh38-based Minigraph-Cactus graph. Variant sites and genotypes were obtained using the PanGenie decompose pipeline (https://github.com/eblerjana/genotyping-pipelines/tree/a7af349abd8fa4d2181b6698a1cf3161588a375f/prepare-vcf-MC), which takes as input a VCF generated using `vg deconstruct` by HPRC2 (https://s3-us-west-2.amazonaws.com/human-pangenomics/index.html?prefix=pangenomes/scratch/2025_02_28_minigraph_cactus/hprc-v2.0-mc-grch38/hprc-v2.0-mc-grch38.vcf.gz. As a preprocessing step, variants with reference allele length exceeding 100 kb were excluded. We disabled the PanGenie pipeline filter that removes bubbles for which more than 20% of the haplotypes carry a missing allele and retained all all variants regardless of missing allele frequency. We then extracted variants of 201 HPRC samples and removed variants with MAS count below 10. We treated the resulting variant set as a truth set, which represents all variants passing the allele count filter across these 201 samples.

#### Variant annotation and classification

We classified each variant by variant type (SNP, small indel, intermediate sized indel and SV) and allelic status (biallelic or multiallelic) based on the maximum length of the reference and alternative alleles after normalization: alleles with ref and alt allele length exact 1 bp were classified as SNPs; alleles with maximum lengths of 2-9 bp as small indels; alleles with maximum lengths of 10-49 bp as intermediate sized indels; alleles with maximum lengths ≥50 bp as SVs. Multiallelic loci were identified by merging normalized records at the same locus using `bcftools norm -m +any` (v1.21). The total number of alleles at each locus was then determined from the REF and ALT fields. Loci with two alleles were classified as biallelic, while loci with three or more alleles were classified as multiallelic. We classified each graph edge in EdgeDepth based on the variant alleles it mapped to. Each allele was first assigned an allele-level type based on the criteria above. Because a graph edge can map to multiple variant alleles, we assigned one summary variant type to each edge using the following priority order, SNP > small indel > intermediate sized indel > SV. An edge was assigned the type of the highest-priority allele among all alleles mapped to it. Edges were classified as biallelic only if they mapped to a single allele at a biallelic locus. All other edges, including edges mapping to multiple alleles or to alleles at multiallelic loci, were classified as multiallelic. Variants in the danbing-tk callset, which only represents tandem repeats, were classified as multiallelic SVs. This classification was used as a summary annotation for downstream enrichment and interpretation analyses.

To annotate the genomic context of each variant, we first identified a GRCh38 interval for each variant. GRCh38 intervals for edges were identified based on the variant alleles mapped to that edge. For edges mapped to multiple alleles, we selected the allele with the shortest GRCh38 span to provide the most specific reference interval for that edge. We then intersected these intervals with genomic annotation tracks using `bedtools intersect` (v2.30.0)^46^. Annotation tracks included low-mappability and segmental duplication regions, tandem repeats ≥101 bp, other difficult genomic regions, and easy regions for variant calling. The first 3 tracks are defined by GIAB genomic stratifications^47^, and easy regions are defined by panmask (pm151b v2)^48^. Because a single variant interval could overlap multiple annotations, we assigned one summary genomic region category to each variant using the following priority order: low-mappability and segmental duplication regions, tandem repeats, other difficult regions, and easy regions. Variant intervals that did not overlap any of these annotation tracks were also assigned to the “other difficult regions” category. This genomic region annotation was used for all downstream enrichment and interpretation analyses.

### Cross-callset variant comparisons

#### Evaluating the performance of short-read variant calling methods as compared to the HPRC2 graph

We normalized graph based variants by splitting multiallelic to biallelic and left-aligning using `bcftools norm` (v1.21). We then compared 1KG, PanGenie, and EdgeDepth callsets against the baseline HPRC2 graph-based variant callset. For each pairwise comparison, we used from `RTG vcfeval`(v3.13)^49^ to deal with possible differences in complex variant representation, which limits to variants with a maximum allele length of 1,000 bp, using `-ref-overlap` to handle overlapping variants and `--sample ALT` to compare genotypes derived from the ALT alleles, using the following command structure:

`rtg RTG_MEM=60G vcfeval -b <graph_vcf> -c <call_vcf> -o <output> -t <GRCh38_SDF> --sample ALT --output-mode annotate --all-records --ref-overlap --no-roc`

Since vcfeval skips evaluation in regions of high complexity, such as some short tandem repeats, we added a patch for these regions. Specifically, if a variant in the baseline callset could not be evaluated by vcfeval, it was considered captured if a variant in the comparison callset shared the same chromosomal position, reference allele, and alternative allele. A baseline variant was ultimately classified as captured if it was called as a true positive (TP) by vcfeval, or if it matched a variant in the comparison callset by exact position and allele sequences under the patch criterion.

EdgeDepth results were directly compared to graph-derived variants using the graph structure. A graph-based variant has a corresponding alternative path record in the INFO/AT field in the VCF file, and it is considered captured if any edges in this path are represented in EdgeDepth callset.

#### Evaluating the performance of the PanGenie callset as compared to the HPRC2 T2T-CHM13 graph

Because 2.8% of PanGenie variants failed to lift over from T2T-CHM13 to GRCh38, reducing the capture rate of graph-based variants by PanGenie, we additionally evaluated PanGenie’s capture rate directly in T2T-CHM13 coordinates using the 201 HPRC samples. We compared the original T2T-CHM13 coordinate PanGenie callset with a pangenome graph-based variant callset derived from the HPRC2 T2T-CHM13 Minigraph-Cactus graph. The T2T-CHM13-based graph variant callset was generated using the same procedure as described for the GRCh38-based graph. We removed variants with a MAS count below 10 from both the T2T-CHM13 graph-based and PanGenie callsets. We then compared the two callsets based on graph structure. A graph-based variant was considered captured by PanGenie if its alternative traversal contained the alternative traversal of a PanGenie variant. This T2T-CHM13 coordinate comparison showed a higher PanGenie capture rate of graph variants than the GRCh38-based comparison, with PanGenie capturing 85.5% of variants represented in the T2T-CHM13-based graph as compared to 79% with the GRCh38-based graph.

#### Differences in variant representation between EdgeDepth and graph-based callsets

EdgeDepth and graph-based callsets differ in how variants are represented, which accounts for differences in variant counts even when both are derived from the same graph. EdgeDepth enumerates all possible alleles represented by the graph, resulting in a more complete decomposition of graph variation. In contrast, graph variants are derived from a multiallelic VCF generated by PanGenie decomposition pipeline, which represents nested variants as observed in the input haplotype assemblies and may not capture every possible allele combination. As a result, EdgeDepth tends to report a larger number of variants, particularly at complex and multiallelic loci.

### eQTL mapping in the 201 HPRC samples

#### Construction of a Unified Transcriptome Annotation

PacBio Kinnex long-read RNA-sequencing data from 206 HPRC2 samples were analyzed to construct a unified transcriptome annotation. For each sample, full-length non-concatemer (FLNC) BAM files generated from multiple sequencing runs were concatenated into a single FLNC BAM file. FLNC reads were converted to FASTQ format and aligned to the GRCh38 reference genome using minimap2 (v2.30)^50^ in high-quality spliced-alignment mode (`-ax splice:hq -uf`), with splice junctions from the GENCODE v48 comprehensive annotation supplied through the `--junc-bed` option after conversion to BED12 format using `paftools.js gff2bed`.

To construct a unified transcriptome annotation across all samples, aligned BAM files from all samples were merged and jointly analyzed using IsoQuant (v3.8.0)^51^, rather than constructing transcript models independently for each sample and subsequently merging them. This approach enhances sensitivity for detecting lowly expressed transcripts. To improve computational efficiency, the merged BAM file was partitioned by chromosome and processed independently. For chromosome 14, only reads mapping to positions 1–104,474,600 were used during transcript model construction because of the extremely high read depth in the immunoglobulin heavy-chain (IGH) region. This depth arises because the samples are lymphoblastoid cell lines derived from B cells with strong IGH expression. The current model-construction algorithm cannot efficiently process this region. However, all chromosome 14 reads were retained for subsequent analyses. The per-chromosome extended GTF files generated by IsoQuant, consisting of the complete reference annotation together with all discovered novel transcripts, were concatenated to produce a single extended GTF file.

Aligned BAM files from each sample were subsequently reanalyzed with IsoQuant using the extended GTF file as the reference annotation and with transcript model construction disabled (`--no_model_construction`). In this mode, IsoQuant performed read-to-transcript assignment and transcript abundance quantification based on the similarity of splice-junction and exon structures between reads and annotated transcripts. Transcript-level count tables generated for individual samples were merged to create a cohort-wide transcript count matrix. Unexpressed transcripts, defined as transcripts with zero counts across all samples, were removed from both the extended GTF file and the transcript count matrix.

Transcripts retained after removal of unexpressed transcripts were subsequently evaluated using SQANTI3 (v5.5.1)^52^, which employs a random forest classifier to distinguish artifact transcripts from true transcripts. The classifier incorporated transcript support derived from the cohort-wide count matrix, splice-junction support from recount3 (srav3h; approximately 228 million junctions from approximately 316,000 public human RNA-seq samples in the Sequence Read Archive)^53^ accessed through Snaptron^54^, transcription start site support from refTSS (v4.1)^55^, polyadenylation motifs, and polyadenylation peak support from PolyASite (v2.0)^56^. Novel transcripts passing SQANTI3 quality-control filtering were combined with GENCODE v48 reference transcripts to generate the unified transcriptome annotation.

#### Transcript Abundance Estimation

Transcript sequences were extracted from the unified transcriptome annotation using GffRead (v0.12.7)^57^. FLNC reads from each sample were aligned to the transcript sequences using minimap2 (v2.30) with parameters `-ax map-hifi --eqx -N 100`, and the resulting alignments were used for transcript abundance estimation with oarfish (v0.9.0)^58^ using parameters `--min-aligned-fraction 0.8 --strand-filter fw --model-coverage`. Transcript abundance estimates from individual samples were merged to generate a transcript-level expression matrix. A gene-level expression matrix was generated by summing transcript abundances for transcripts assigned to the same gene. These matrices were used for eQTL analyses. The transcriptome construction and quantification workflows were implemented using Nextflow^59^ and are publicly available at https://github.com/wwliao/hprc_release2_kinnex_analysis.

#### Gene expression quantification and covariate selection

For the cohort of 201 HPRC samples, we extracted gene expression levels quantified from PacBio Kinnex long-read sequencing data as described above. We followed AFGR study gene filtering criteria and covariate identification procedures^17^. We removed lowly expressed genes, defined as genes with a mean raw count fewer than five reads across samples. We then applied variance-stabilizing transformation to retained gene counts using DESeq2^41^, followed by standardization to zero mean and unit variance. Covariates for eQTL analyses include genotype principal components (PCs), hidden factors, one-hot encoded generator facility (Rockefeller, UW, WashU) and standardized median read length. Genotype PCs were computed using SNPRelate (v1.32.2)^60,61^ on all variants from these samples included in the 1KG callset. The number of PCs was determined by selecting the minimum number of PCs needed to cumulatively explain at least 5% of genetic variation. Hidden confounding factors were identified using SVA (v3.46.0)^62^, retaining the maximum number of surrogate variables. This resulted in 2 PCs and 16 surrogate variables.

#### cis-eQTL mapping using graph variants

For each gene, we considered all graph variants passing filters within 1 Mb of the transcription start site (TSS). We computed nominal eQTL p-value for each variant-gene pair and permutation-based p-values for the lead eQTL variant per gene using tensorQTL (v1.0.8)^21^, with 10,000 permutations (random seed: 1234). Gene-level significance thresholds (q-value and nominal p-value threshold) were estimated using a beta distribution approximation to the permutation p-value distribution. eGenes were defined as genes with a lead variant q-value < 0.05, and significant variant-gene pairs were defined as those with a nominal p-value below nominal p-value threshold of the gene. For variants with identical nominal p-values (ties) as lead eQTL markers, we set `random_tiebreak=True` in tensorQTL. We identified lead eQTL markers for each eGene and calculated the lead variant capture rates by the 1KG, PanGenie, and EdgeDepth callsets.

#### Comparison of eQTL discovery between graph variants and 1KG variants

To compare eQTL discovery between graph-based variants and 1KG variants, we performed a joint cis-eQTL analysis by pooling variants from the 1KG and graph-based callsets, using the same eQTL analysis settings described in the cis-eQTL mapping using graph variants section. A callset was considered to identify an eGene if it contributed a variant with a nominal p-value below the gene-level p-value significance threshold. We defined graph-specific eQTL variants as the lead markers of eGenes identified only by graph-based variants, that is, eGenes for which no 1KG variants reached the gene-level significance threshold. We then calculated the capture rates of these graph-specific lead variants by the 1KG, PanGenie, and EdgeDepth callsets.

### eQTL mapping in the 430 AFGR samples

#### Variant matching and rescue across callsets

Because the same variant might be represented redundantly across multiple callsets. We identified matched variants among 1KG, PanGenie, and EdgeDepth prior to minor allele sample (MAS) filtering. We determined variant matching either by true-positive matches identified by `RTG vcfeval` (v3.13)^49^ or by identical chromosome, position, reference allele and alternative allele after left-alignment and normalization with `bcftools norm`. Because MAS filtering for EdgeDepth is derived from GMM model fitting rather than discrete genotype counts, which was used for 1KG and PanGenie, the filtering could be slightly different across the callsets. To reduce the possibility of callset-specific eQTL signals arising from a variant being retained in one callset but filtered in another, we applied a rescue step. After applying the primary MAS < 10 filter, a variant was rescued and included in all downstream eQTL analyses if it had MAS ≥ 5 and its matched variant in at least one other callset had MAS ≥ 10. The rescue step is applied symmetrically across all three callsets.

#### Expression normalization and eQTL mapping

Gene raw counts from LCL RNA-seq data were quantified using GENCODE annotation (v27) by the AFGR project^17^. Expression normalization, covariate selection (16 PCs, 24 surrogate variables), cis-eQTL mapping, and lead-marker identification were done exactly as described above for the PacBio Kinnex analysis of the HPRC samples. To ensure fair comparison of eQTL discovery across callsets, we quantified tie cases between 1KG and PanGenie based on nominal p-value, both of which use discrete genotypes, and evenly split tied lead variants between the two callsets. EdgeDepth and danbing-tk use continuous dosage values, which do not produce ties and were unaffected by this adjustment. We performed eQTL analyses using the same expression matrix, covariates, parameters and thresholds for (1) each variant callset separately, (2) a joint callset constructed by pooling variants from all callsets, and (3) the merged HPRC2+1KG callset representing unique variants across all callsets after redundancy removal. Details of each analysis are described in the following sections.

#### Separate eQTL mapping for each individual callset

To assess the performance of each variant callset in eQTL discovery, we performed cis-eQTL mapping using variants of 430 samples from four callsets separately: 1KG, PanGenie, EdgeDepth and danbing-tk. For each callset, we performed eQTL analysis in 430 samples and recorded the number of eGenes identified and classified the lead eQTL marker by variant type using the annotation procedure as described in the “Variant annotation and classification section”. We compared the number of eGenes and the distribution of lead-marker types across callsets to evaluate differences in the genetic variation captured by each approach.

#### Joint eQTL mapping and callset-specific eGene discovery

To enable direct comparison of eQTL discovery across callsets, we performed a joint eQTL analysis using all variants of 430 samples pooled from 4 callsets (1KG, PanGenie, EdgeDepth, and danbing-tk). Within this joint analysis, an eGene was identified significantly by a given callset if that callset contributed at least one variant with a nominal p-value below the gene-specific nominal p-value threshold. We then counted the number of eGenes identified exclusively by pangenome-based callsets (PanGenie, EdgeDepth, and danbing-tk), i.e., eGenes for which no 1KG variant reached nominal significance.

#### Quantification of pangenome-driven eQTL signal improvement in the joint analysis

To quantify the improvement in association signal from pangenome-based callsets relative to 1KG in the joint eQTL mapping experiment, we computed the percent increase in -log10(nominal p-value) for the most significant variant across all pangenome callsets relative to the most significant variant in the 1KG callset, for each eGene. We then counted eGenes that have strong signal improvement (>50% increase in -log10p-value) and moderate signal improvement (>20% increase in -log10p-value). Among the eGenes with strong signal improvement and with moderate improvement, we annotated the lead marker by variant type and determined whether the improvement was driven by PanGenie, EdgeDepth, and/or danbing-tk by whether each callset report any variant with corresponding percent of improvement.

#### Classification of mechanisms underlying improved lead variants in the joint eQTL analysis

To investigate whether the association signal improvement from pangenome is attributed to novel variants missed by 1KG callset or to improved genotyping accuracy of variants present in 1KG callset, we classified the improved eGenes into 7 mutually exclusive categories: (1) immunoglobulin (IG) gene, (2) tandem repeat, (3) SNP in 1KG, (4) SNP not in 1KG, (5) indel in 1KG, (6) indel not in 1KG and (7) SV. IG genes were treated as a separate category because in this study we used LCLs, in which IG genes are highly expressed and eQTL signals at IG loci should be interpreted cautiously due to somatic rearrangement and potential read mapping ambiguity in these regions. We included the tandem repeat category because GATK is known to struggle with such variants. A lead variant was classified as a tandem repeat if it met either of the following criteria: i) it was in a homopolymer sequence, defined as the inserted or deleted sequence consists a single nucleotide, and that nucleotide is extended in an uninterrupted run of ≥5 bp at the variant position in the GRCh38 reference; or ii) the variant position overlaps a simple tandem repeat located by Tandem Repeats Finder (TRF) ^63^ and the repeat unit sequence comprised >50% of the inserted or deleted allele sequence. For variants not classified as tandem repeats, we categorized lead variants by allele length: SNPs (ref and alt alleles both equal to 1 bp), indels (1 < max(len(ref), len(alt)) < 50 bp), and SVs (max(len(ref), len(alt)) ≥ 50 bp). Each size category was further divided by whether the variant was present in the 1KG callset, as described by the criteria in the variant matching section.

#### Identification and visualization of noteworthy eQTL examples

To identify pangenome-driven eQTL improvement in medically relevant examples, we intersected the 185 eGenes with strong signal improvement (>50% increase in -log10p-value) with a curated set of medically relevant genes ^22^. We selected 4 representative genes of different variant types and improvement reasons and visualized them in detail: *MINPP1*, *SURF1*, *TRIB3*, *CBS*. For each example, pairwise genotype correlation between lead eQTL variant and other variants was quantified as r^2^ derived from Pearson correlation. To visualize the correlation between gene expression and the lead marker genotype, we projected EdgeDepth dosage values onto discrete genotype groups using the genome STRiP constrained GMM described in the “Selection of variable edges section”. For one gene (*TRIB3*), the GMM did not yield well-separated genotype clusters based on visual inspection. In this case, we plotted correlation between gene expression and binned normalized EdgeDepth intervals (0-15, 15-30, 30-45). We visualized the pangenome graph structure at each lead variant locus by extracting the corresponding subgraph using gfabase (v0.6.0, https://github.com/mlin/gfabase) and plotting it by Bandage (v0.8.1) ^64^. In the Bandage plot, reference nodes are colored in light blue and non-reference nodes in dark blue, with the lead edge indicated by an arrow.

#### Head-to-head comparison of eQTL discovery across 1KG, PanGenie, and EdgeDepth callsets

To evaluate the effect of variant genotype quality on eQTL signal strength, we performed pairwise comparisons among the 1KG, PanGenie, and EdgeDepth callsets. For each pair (1KG - PanGenie, 1KG - EdgeDepth, and PanGenie - EdgeDepth), we extracted variants shared between the two callsets and constructed a joint callset in which each variant was represented by both callsets simultaneously. Restricting analysis to shared variants ensures that any difference in eQTL discovery reflects genotype quality rather than variant inclusion. For each pairwise joint callset, we identified eGenes and compared the number of lead variants identified by each callset. We also compared the -log10(nominal p-value) of the most significant variant from each callset across all eGenes, and visualized them as scatter plots with one point per eGene.

### eQTL mapping using the merged “HPRC2+1KG” callset

#### Merged callset generation

We constructed a merged variant callset to represent all unique variants across the 430 AFGR samples, integrating 4 callsets: 1KG, PanGenie, EdgeDepth, and danbing-tk. We removed variants reported redundantly across the 1KG, PanGenie, and EdgeDepth callsets, based on variant matching and genotype correlation. The method for identifying matched variants is described in the “Variant matching and rescue across callsets section”. When a variant matched multiple variants in another callset, we selected a single variant with the highest squared Pearson correlation coefficient (r^2^) as the matched pair. We then calculated the r^2^ between variant match pair genotypes (for 1KG and PanGenie) or dosages (for EdgeDepth) across 430 samples and only treated variant match pairs with r^2^≥ 0.5 as true redundant variants. The reason for this is that we noticed instances where seemingly matched variants had very different genotype information, suggesting they were in fact representing different variants. The exception to this process was danbing-tk variants, which represent the dosage of each tandem repeat and were not well captured by the other three callsets and therefore retained in full. We then constructed the merged callset by removing redundant variants and retaining a single representation per variant according to the priority order of 1KG, PanGenie, EdgeDepth, and danbing-tk, with variants represented in a higher-priority callset removed from lower-priority callsets. The merged callset represented 1KG calls supplemented by pangenome-specific variants. We then performed cis-eQTL mapping using this merged callset in the 430 AFGR samples (as described in “Expression normalization and eQTL mapping section”), reporting eGenes, permutation p-value for lead makers per eGene, and nominal p-value for all variant - gene pairs.

#### Fine mapping of causal variants

We identified independent signals for each eGene from the merged callset using Sum of Single Effects (SuSiE) regression (v0.12.35)^65,66^. For each eGene, we extracted all variants within 1 Mb window of the TSS, regressed covariates out from both the genotype matrix and gene expression values, and applied the `susie` function with a maximum of 10 non-zero effects. We reported 95% credible set (CS) for each independent signal. We then quantified the contribution of pangenome-specific variants to each CS by summing the per signal posterior inclusion probabilities (PIPs) of all pangenome variants within that CS.

To assess the added value of pangenome-specific variants in fine-mapping, we repeated the analysis using 1KG variants alone, and compared results between the 1KG alone and merged callset fine-mapping tracing the PIP shift from 1KG to pangenome specific variants for each eGene. We first assigned each CS to one of five categories based on its pangenome variant contribution and matching across the two fine-mapping runs: (1) pangenome-dominant, where pangenome variants accounted for more than half the total CS PIP (pangenome PIP > 0.5); (2) pangenome-contributing and matched, where pangenome variants contributed moderately (0.1 < pangenome PIP ≤ 0.5) and the CS shared at least one variant with a CS from the 1KG alone run; (3) stable and matched, where the CS was concordant across both runs and pangenome variants did not contribute meaningfully or at all (pangenome PIP ≤ 0.1); (4) lost, where the CS was present in the 1KG alone callset run but had no matching CS in the merged run; and (5) novel, where the CS was present only in the merged callset run. We then assigned each eGene to one of four gene-level categories based on the composition of its CS categories: (1) pangenome refined signal, with at least one CS in category 1, indicating pangenome variants improved signal discovery; (2) pangenome expanded signal, with no CS in category 1 but at least one CS in category 2, indicating pangenome variants added additional candidates to the signal; (3) stable signal, with all CSs in category 3, indicating fine-mapping was unaffected by the addition of pangenome-specific variants; and (4) lost signal, with no CSs in categories 1 or 2, but at least one CS in category 4, indicating that inclusion of pangenome-specific variants diffused previously identified signals. We counted the number of eGenes in each category.

#### Gene expression prediction

We predicted gene expression using variants within 1 Mb of the TSS using 1KG callset and the merged HPRC2+1KG callset, and compared prediction performance. The target gene set comprised the union of eGenes identified across callsets. For each callset, we regressed covariates out of both the genotype matrix and gene expression values and scaled genotype and gene expression prior to model fitting. We trained gene expression prediction models using 5 methods: Lasso, ridge regression, elastic net, top-1 (single best variant), and SuSiE, and evaluated them by five-fold cross-validation. Elastic net and top-1 implementations were extracted from the FUSION package^28^. Although FUSION also includes LASSO and ridge regression, its implementations are restricted to discrete genotypes and are therefore incompatible with the continuous dosage values used by EdgeDepth. We therefore reimplemented LASSO and ridge regression using glmnet (v4.1.8)^67–69^. SuSiE-based prediction was performed using susieR (v0.12.35)^65,66^. For each gene and callset, we assessed prediction accuracy by fitting a linear regression of predicted expression on observed expression and computing the adjusted r^2^ as implemented in the FUSION package. We then selected the highest adjusted r^2^ across the 5 methods as the prediction result for each gene and each callset.

### Comparison of eQTLs and GWAS loci

#### Colocalization analysis

We performed eQTL-GWAS colocalization for 812 eQTLs identified by joint eQTL analysis in which pangenome variants showed signal improvement over 1KG variants (defined as >20% increase in -log10p-value, see “Quantification of pangenome-driven eQTL signal improvement” section). We selected 41 GWAS studies from GWAS catalog^70^ with harmonized summary statistics, including blood biomarkers (e.g. lymphocyte count), common complex disease (e.g. Type 2 diabetes), and conditions with etiology related to immune function (e.g. Crohn’s disease) as in the AFGR study^17^. We identified the candidate eQTL-GWAS pairs by requiring at least one GWAS signal with p-value < 5 x 10^-8^ within 100 kb of the eQTL lead markers. We then performed colocalization using coloc (v5.2.3)^27,31,71^ under the assumption of a single causal variant affecting the trait. We restricted the input summary statistics to 1KG variants within 500 kb of the eQTL lead marker that were present in both the eQTL and GWAS studies. Variant matching across the studies was based on identical chromosome, position, reference allele, and alternative allele. We identified ten eQTL-GWAS pairs with strong evidence of colocalization, defined as those with posterior probability H4 ≥ 0.8, where H4 represents the hypothesis that both traits have genetic association in the region and share a single causal variant, and moderate evidence of colocalization as those with 0.5 ≤ H4 < 0.8.

#### Visualization of colocalized eQTL-GWAS loci

We visualized the 10 loci using LocusZoom^72^ plots, expression genotype correlation, and pangenome graph structure. For each locus, we quantified LD between the lead variant and all other variants in the window using the squared Pearson correlation coefficient (r^2^). We visualized each loci by two LocusZoom plots colored by LD with lead variants and by variant callset respectively. We also visualized the correlation between gene expression and lead marker genotype. Because EdgeDepth reports continuous depth rather than discrete genotypes, we assigned genotype classes to each sample by fitting the GenomeSTRiP constrained GMM (as described in the “Selection of variable edges section”) to each edge and assigning each sample to one genotype cluster based on its probability in that distribution. We then visualized the pangenome graph structure at each lead variant locus by extracting the surrounding subgraph from HPRC2 Minigraph-Cactus graph (GRCh38 version) using gfabase (v0.6.0, https://github.com/mlin/gfabase) and plotting it by Bandage (v0.8.1)^64^. In the Bandage plot, reference nodes are colored red and the lead edge and its adjacent non-reference nodes are colored blue.

#### Analysis of the *GBAP1* locus

We performed a detailed analysis of the *GBAP1* locus to characterize structural alleles and relate the lead eQTL edge to *GBAP1* copy number variation. *GBAP1* is a pseudogene of *GBA1* with high sequence identity. To locate all copies of *GBAP1* and *GBA1* in the pangenome graph, we extracted the subgraph spanning chr1:155,212,662-155,245,699 from the HPRC2 Minigraph-Cactus graph (GRCh38 version) using gfabase, where this genomic interval was selected to encompass all gene copies. We then aligned the *GBAP1* and *GBA1* reference sequences (GENCODE v27^73^) to the subgraph using GraphAligner (v1.0.13)^74^ to identify the graph nodes and bubbles corresponding to each gene copy. We traced each HPRC haplotype assembly path (N=464) through these nodes to determine the structural alleles and *GBAP1* copy number per haplotype and calculate the number and frequency of haplotypes carrying each structural allele. We visualized structural haplotypes as linear gene diagrams using gggenes (v0.6.0, https://github.com/wilkox/gggenes), with gene lengths drawn proportional to annotated gene length.

## Data availability

Short-read WGS for 1KG samples were downloaded from the European Nucleotide Archive under accession PRJEB31736: https://www.ebi.ac.uk/ena/data/view/PRJEB31736. RNA-seq data for the AFGR samples were obtained from the dataset published in ref^17^. Long-read PacBio Kinnex RNA-seq data are available from: https://github.com/human-pangenomics/hprc_intermediate_assembly/blob/main/data_tables/sequencing_data/data_kinnex_pre_release.index.csv. The 1KG GATK variant callsets were downloaded from: https://ftp.1000genomes.ebi.ac.uk/vol1/ftp/data_collections/1000G_2504_high_coverage/working/20201028_3202_phased. The 1KG structural variant callset was downloaded from: http://ftp.1000genomes.ebi.ac.uk/vol1/ftp/data_collections/1000G_2504_high_coverage/working/20210124.SV_Illumina_Integration/1KGP_3202.gatksv_svtools_novelins.freeze_V3.wAF.vcf.gz. The PanGenie callset for the 1KG samples was obtained from: https://s3-us-west-2.amazonaws.com/human-pangenomics/pangenomes/scratch/2026_03_30_pangenie/pangenie_all-samples_filtered.vcf.gz. The danbing-tk callset was obtained from: https://sandbox.zenodo.org/records/277309?preview=1&token=eyJhbGciOiJIUzUxMiJ9.eyJpZCI6IjgyNDBiMzE4LTRiOGEtNGYyZS04MzA0LTBmNjI1MGM2NWU4MiIsImRhdGEiOnt9LCJyYW5kb20iOiI0YWQzY2Y1MzFiZWY0YzRkZmExNTFjNmRhMjhjMDMxOSJ9.aLMMXRR9T7ywhYTf9_N46sZ7ppTRVkyvcXCdSlP42xX_9N-lWtOiQ7Xy-LtsZrHGoRT9joycIEISUvUP0h4VYA. GIAB benchmark regions were downloaded from: https://ftp-trace.ncbi.nlm.nih.gov/giab/ftp/release/genome-stratifications/v3.6/GRCh38@all/. Panmask easy regions were downloaded from: https://doi.org/10.5281/zenodo.15683328. The HPRC2 GRCh38 Minigraph-Cactus graph was downloaded from: https://s3-us-west-2.amazonaws.com/human-pangenomics/pangenomes/scratch/2025_02_28_minigraph_cactus/hprc-v2.0-mc-grch38/hprc-v2.0-mc-grch38.gbz. The HPRC2 CHM13 Minigraph-Cactus graph was downloaded from: https://s3-us-west-2.amazonaws.com/human-pangenomics/pangenomes/scratch/2025_02_28_minigraph_cactus/hprc-v2.0-mc-chm13/hprc-v2.0-mc-chm13.gbz. Sample list for the 201 HPRC and 430 AFGR samples, the EdgeDepth callsets for the 430 AFGR samples, the file linking graph edges to corresponding genetic variants, and the eQTL summary statistics, are available from: https://doi.org/10.5281/zenodo.20938527.

## Code availability

The implementation of EdgeDepth pipeline is available at: https://github.com/lushjia/EdgeDepth. Source code for the PacBio Kinnex RNA-seq data analysis pipeline is available at: https://github.com/wwliao/hprc_release2_kinnex_analysis.

## Acknowledgments

This work was funded by NIH/NHGRI awards R01HG013371 (I.M.H. and N.O.S.) and U41HG010972 (I.M.H.). M.J.P.C. and T.-Y.L. were supported by NIH/NHGRI awards R01HG011649 and U01HG010973. T.M. and J.E. were supported by NIH/NHGRI award U01HG013748.

## Author Contributions

S.L. performed all analyses related to the discovery and interpretation of genetically regulated gene expression. W.-W.L. developed the original EdgeDepth method, which was evaluated and refined with S.L. W.-W.L. led the analysis of long-read RNA-seq data. T.-Y.L. and M.J.P.C. provided the danbing-tk callset. J.E. and T.M. provided the PanGenie callset. M.K.D., P.C.G. and S.B.M. provided the AFGR RNA-seq data and advised on the analysis of AFGR data. S.L. and I.M.H. wrote the paper. All authors reviewed the manuscript and provided comments and edits. I.M.H. and N.O.S. conceived the study.

## Supplementary Figures

**Supplementary Figure 1.**
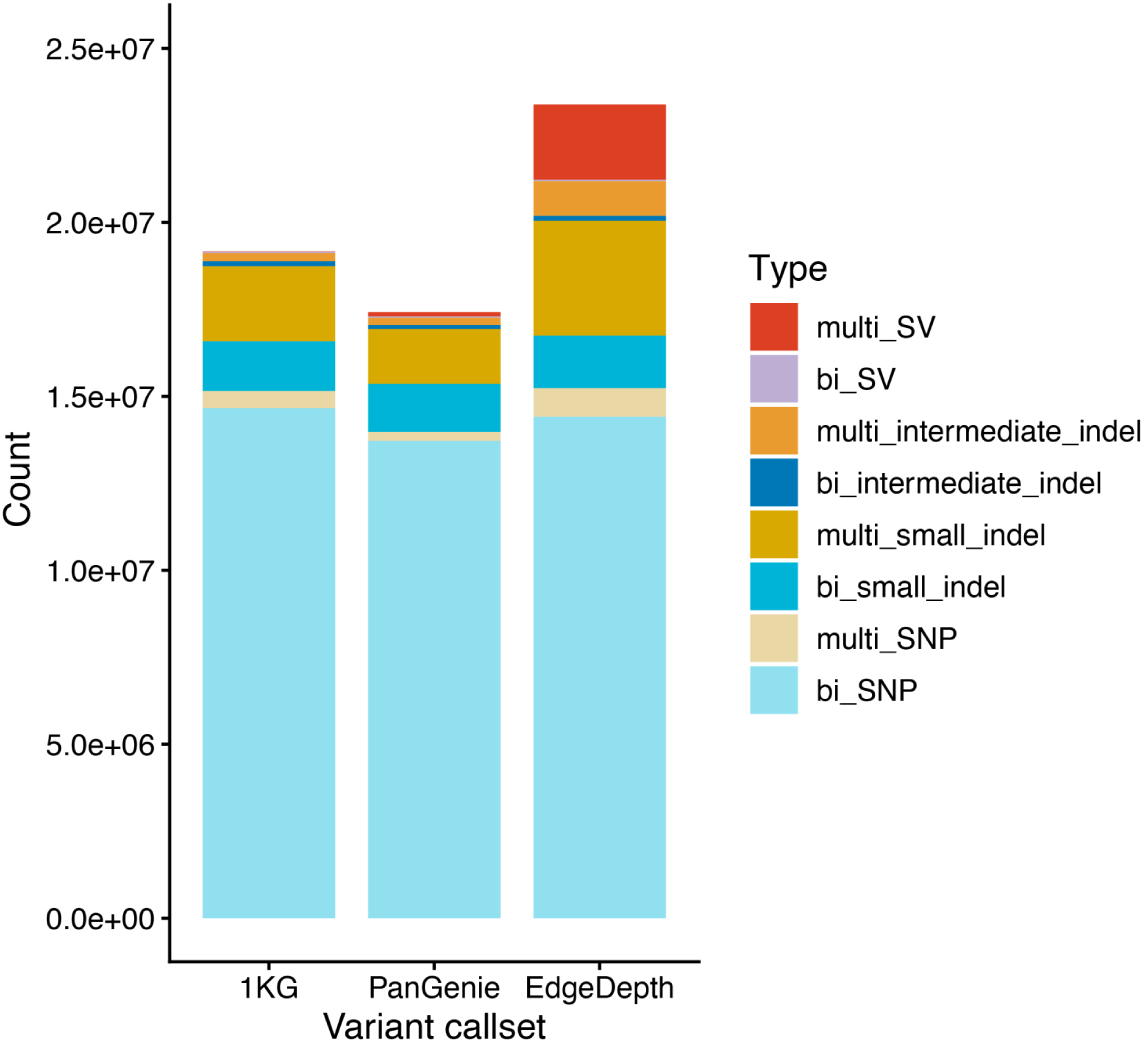
Variant type composition in each callset. Variant counts are shown by type after filtering for variant quality and allele frequency, and after rescue of variants matched across callsets (see Methods). Variant types include biallelic or multiallelic variants (bi_ or multi_) and SNPs (1 bp), small indels (2-9 bp), intermediate sized indels (10-49 bp) and SVs (≥50 bp).

**Supplementary Figure 2.**
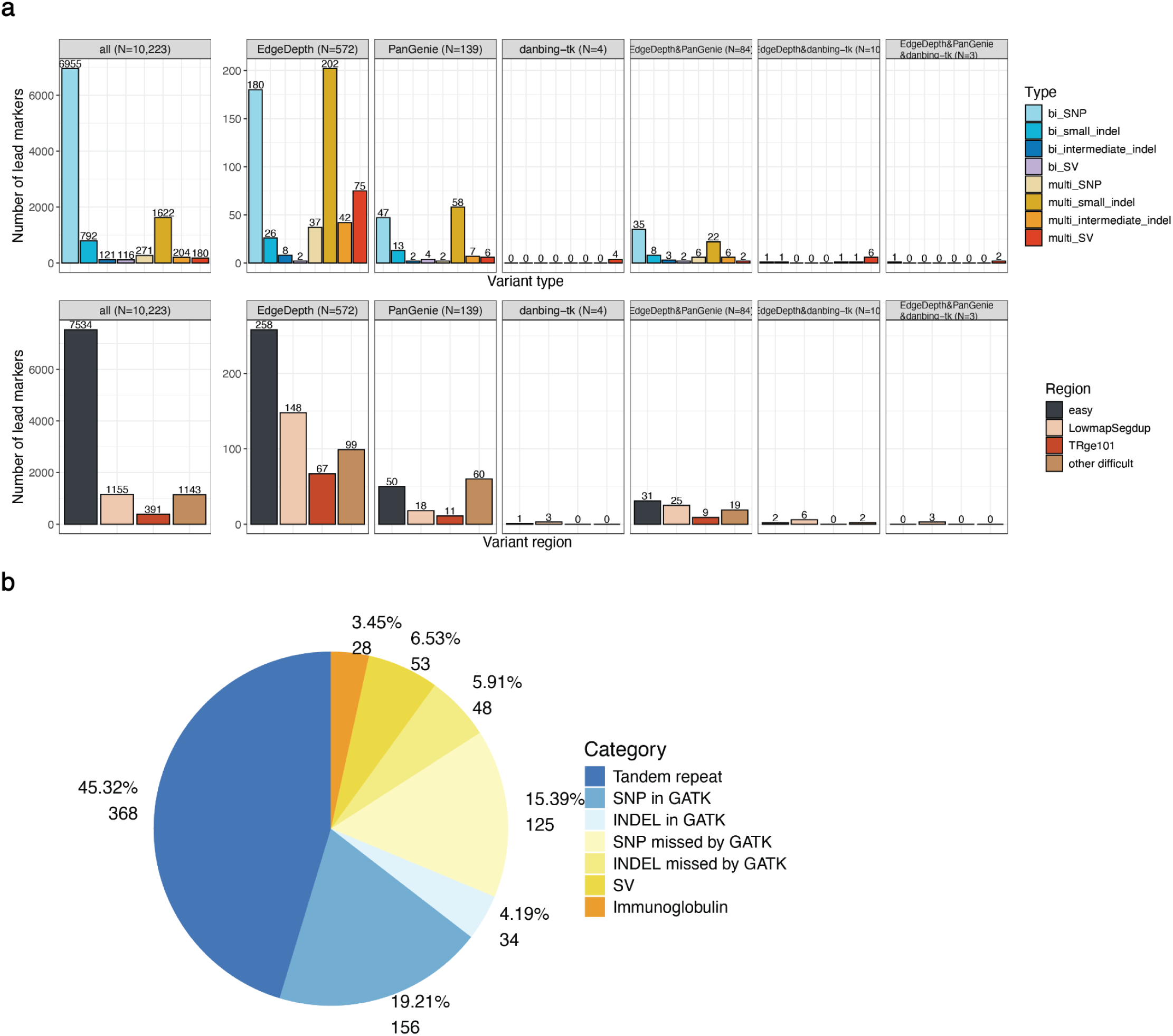
The 812 eGenes with a pangenome power boost. Variant type composition of lead eQTL markers and the underlying mechanisms of the 812 eGenes at which pangenome variants improved the eQTL signal, defined as a >20% increase in -log10(nominal p-value) for the most significant variant across all pangenome callsets relative to the most significant 1KG variant. This figure follows the same format as Fig. 3c,d. **(a)** Composition of lead eQTL markers by variant type (top) and genomic region (bottom). The left panel shows all eGenes from the joint analysis (N = 10,223). The right panels show the 812 improved eGenes split by the driving callset, defined by if the callset reported any variant with >20% of improvement. **(b)** Pie chart of the mechanisms underlying the 812 improved lead variants, including tandem repeat, SNP in 1KG, INDEL in 1KG, SNP missed in 1KG, INDEL missed by 1KG, SV, and immunoglobulin.

**Supplementary Figures 3 and 4.**
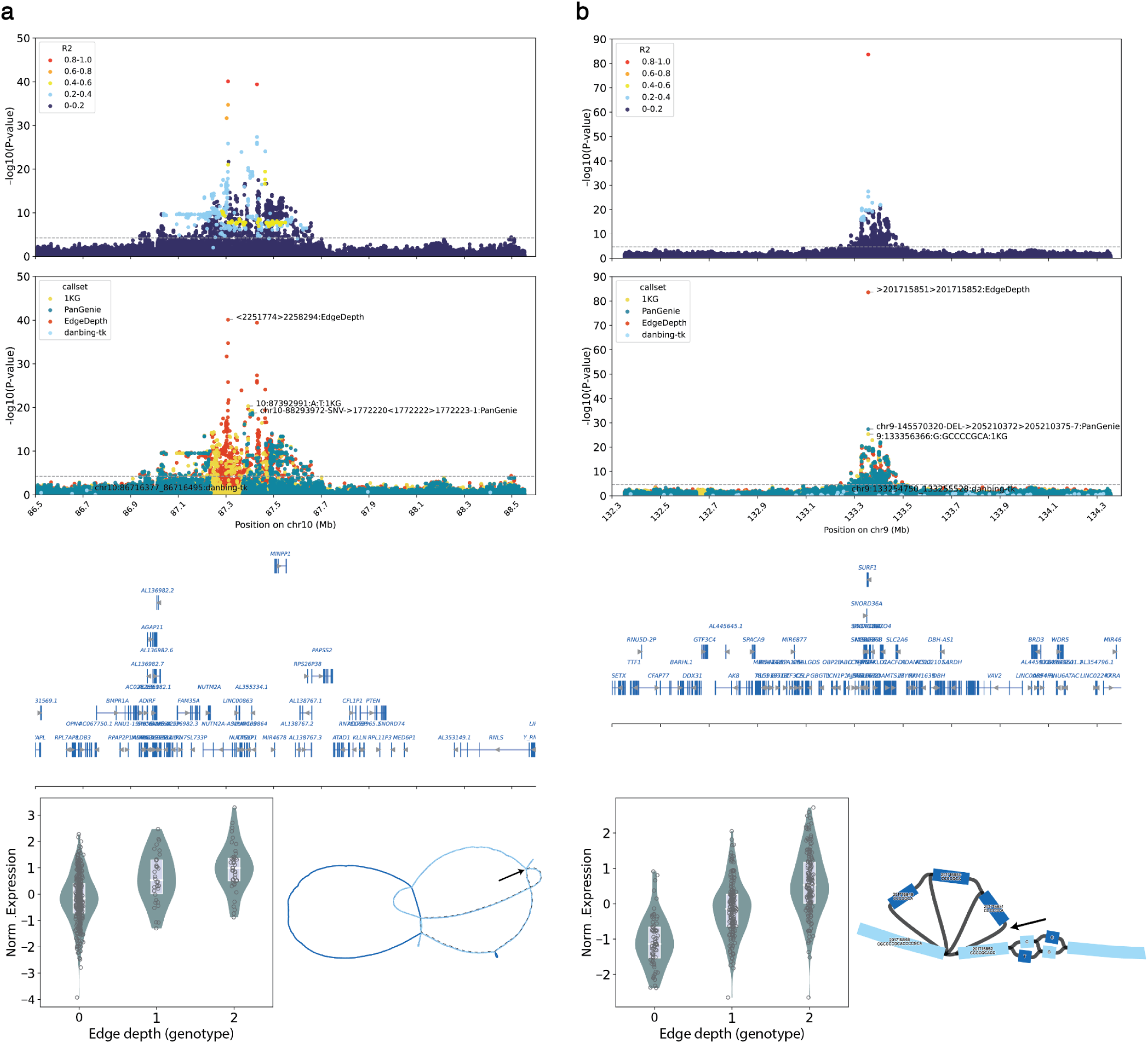
eQTL signals and pangenome graph structures for noteworthy examples. Shown are four representative genes: *MINPP1* (Supplementary Fig. 3a), *SURF1* (Supplementary Fig. 3b), *TRIB3* (Supplementary Fig. 4a), and *CBS* (Supplementary Fig. 4b). Each example is shown in five panels from top to bottom: (1) a LocusZoom plot of association within 1 Mb of the transcription start site (TSS) of the gene color by r^2^ (LD) to the lead eQTL marker; y-axis, −log10(nominal p-value); x-axis, GRCh38 position; the dashed line marks the gene-specific nominal p-value significance threshold; (2) a Locuszoom plot as in (1) colored by callset, with the lead marker of each callset annotated; (3) gene annotations; (4) normalized gene expression versus the lead-marker genotype across 430 AFGR samples. Edge depth genotype is called from fitting a constrained Gaussian mixture model (GMM) from genomeSTRiP to continuous edge depth dosage. Boxes show the median and interquartile range (25th - 75th percentile). Individual samples are shown as points; and (5) a Bandage plot of the pangenome graph structure surrounding the lead variant in the pangenome graph. Reference nodes are colored in light blue and non-reference nodes in dark blue, with the lead edge indicated by an arrow. The dotted line in Supplementary Fig. 3a represents the inversion.

**Supplementary Figure 4.**
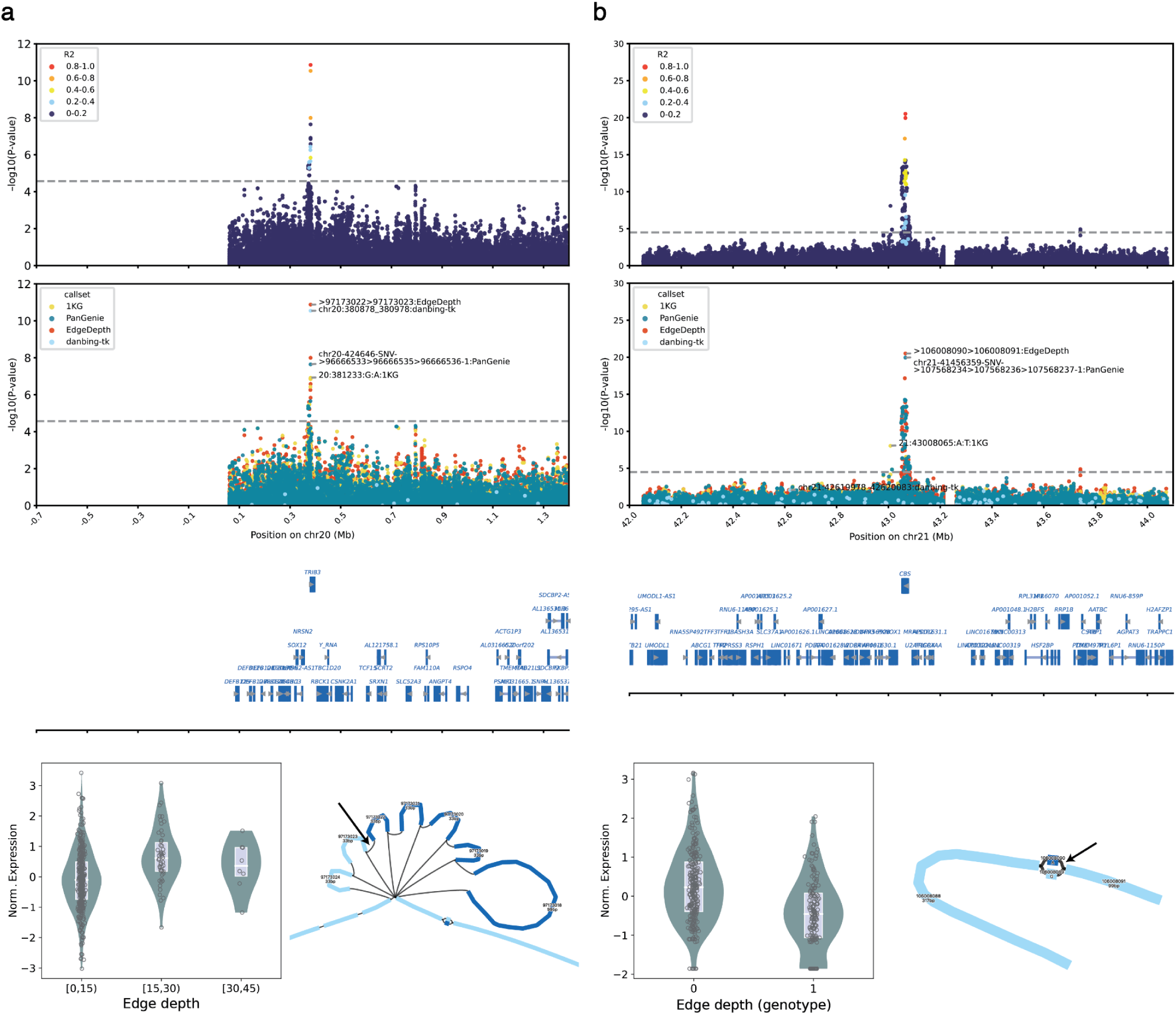
See the legend for Supplementary Figure 3.

**Supplementary Figure 5.**
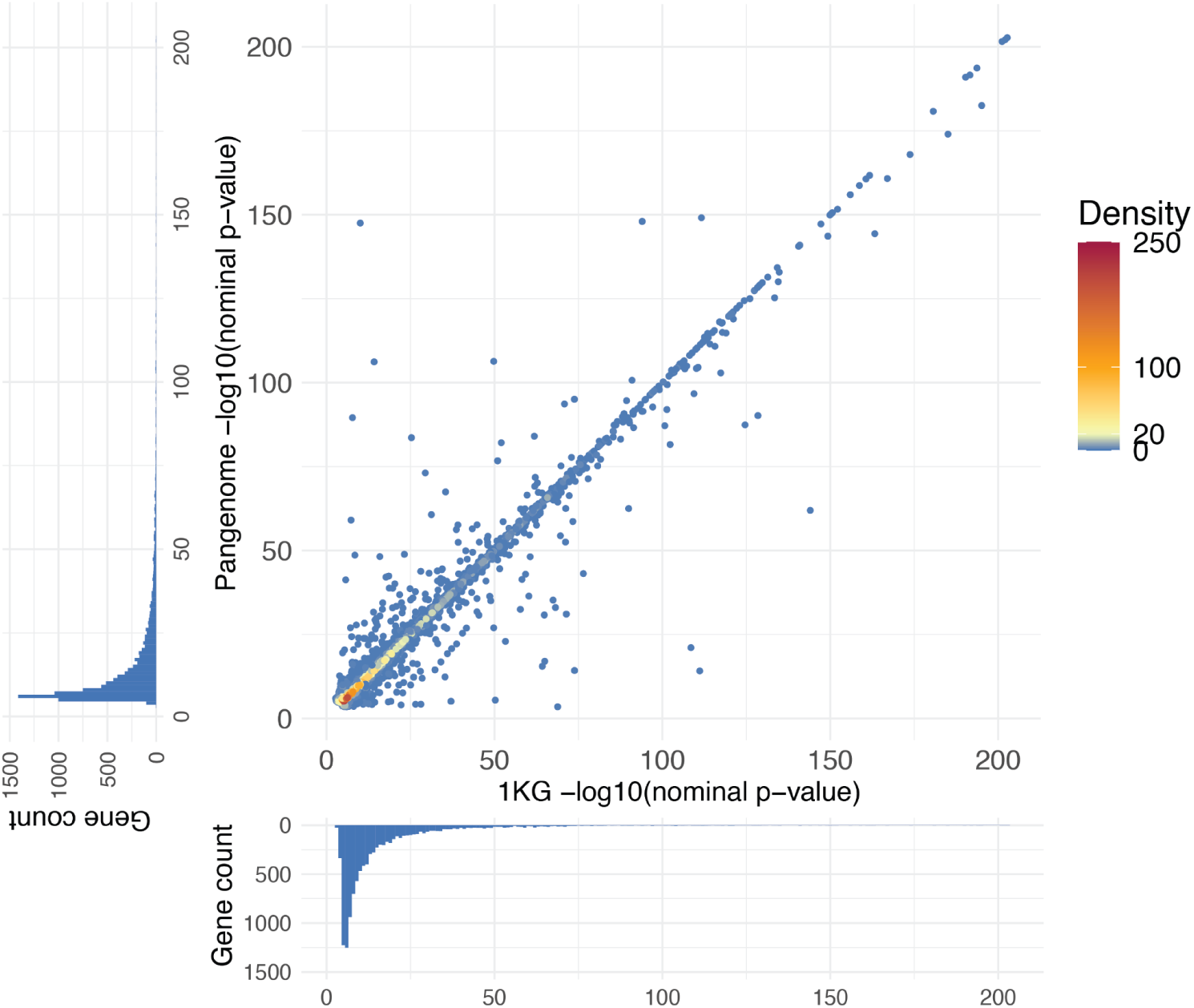
Comparison of eQTL significance in pangenome callsets versus 1KG. Scatter plot comparing the strongest eQTL association signal from pangenome callsets with the strongest signal from the 1KG callset for each eGene in the joint eQTL analysis (N=10,223). For each point (eGene), the y axis shows the strongest -log10(nominal p-value) among variants from pangenome callsets, and the x axis shows the strongest -log10(nominal p-value) among variants from the 1KG callset. Points are colored by local density. Marginal histograms show the distributions of -log10(nominal p-value) for each axis.

**Supplementary Figure 6.**
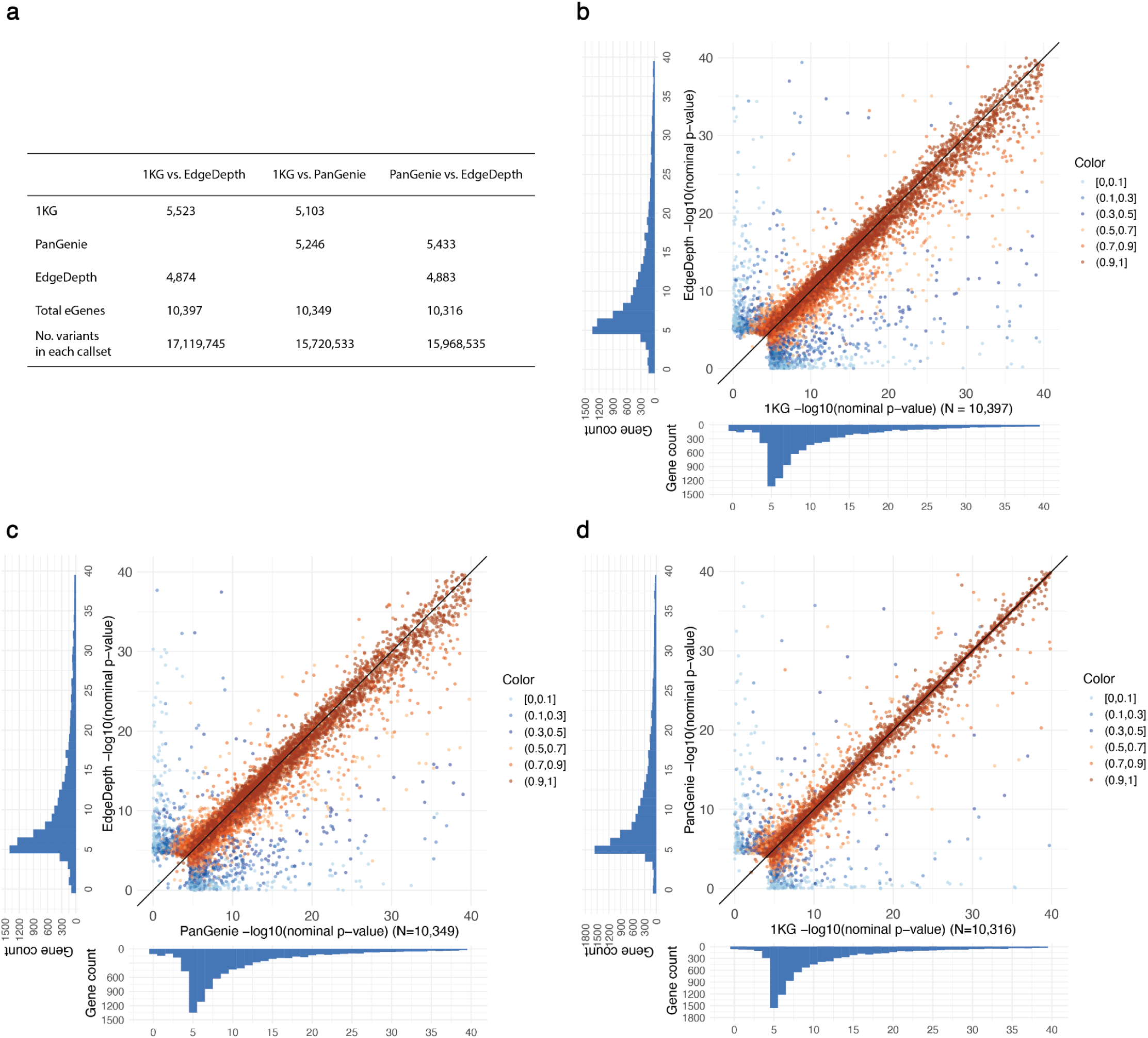
Head-to-head comparison of eQTL significance at shared variants. Pairwise comparisons were performed among the 1KG, PanGenie, and EdgeDepth callsets using variants shared between each pair of callsets. A joint cis-eQTL analysis was performed for each pairwise comparison: 1KG vs. PanGenie, 1KG vs. EdgeDepth, and PanGenie vs. EdgeDepth. **(a)** Summary table showing, for each pairwise comparison, the number of eGenes contributed by each callset, the total number of eGenes identified, and the number of shared variants included in the analysis. **(b-d)** Scatter plots comparing eQTL association signals between each pair of callsets, zoomed to the range 0-40 on both axes. Each point represents the -log10(nominal p-value) of a lead variant from one callset and its corresponding matched variant in the other callset. Points are colored by the correlation between the lead variant and its matched variant. Marginal histograms show the distributions of -log10(nominal p-value) for each axis.

**Supplementary Figure 7.**
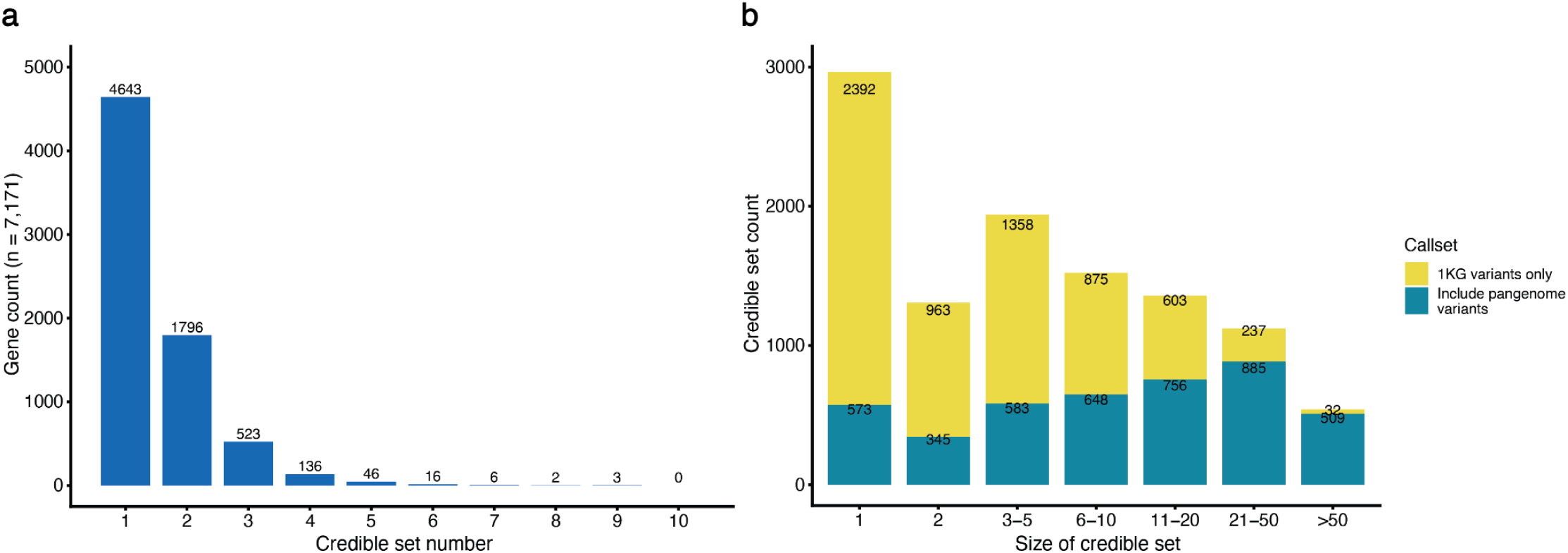
Summary of credible sets (CSs) at fine-mapped eQTL signals from the merged HPRC2+1KG callset. **(a)** Histogram showing the number of credible sets per eGene (N = 7,171). **(b)** Histogram showing the number of variants per credible set. Each bar is colored to indicate how many credible sets of that size range include pangenome-specific variants or 1KG variants only.

**Supplementary Figure 8.**
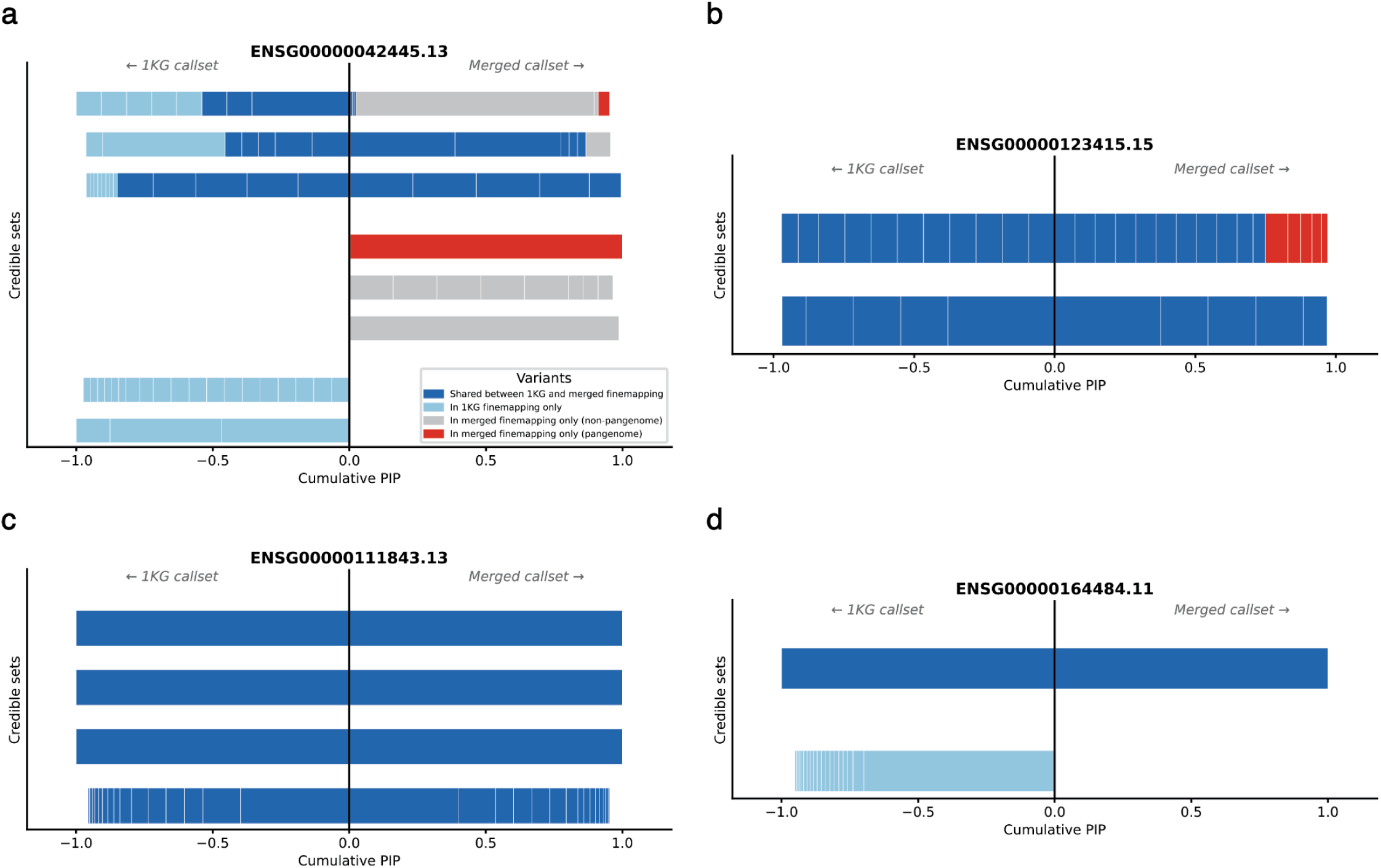
Example fine-mapping results showing effects of pangenome-specific variants. Four representative examples illustrating categories of eGenes according to how posterior inclusion probabilities (PIPs) shift after adding pangenome-specific variants to 1KG for fine-mapping. Categories include pangenome refined signals, pangenome expanded signals, stable signals, and lost signals. **(a)** Example of “Pangenome refined” signals, defined as eGenes with at least one credible set (CS) dominated by pangenome-specific variants (summed pangenome variant PIP > 0.5), indicating that pangenome variants markedly improved signal discovery. Comparison of credible sets identified by fine-mapping using the 1KG callset alone (left) and the merged HPRC2+1KG callset (right). Each row represents one credible set (CS). Aligned rows indicate matched CSs, defined as CSs that share at least one variant across the two fine-mapping runs, whereas unaligned rows indicate CSs without a match in the other run. Variants within each CS are represented by rectangles and colored by matched status across the two runs and pangenome-specific variants. **(b)** Example of “pangenome expanded” signals, defined as eGenes where no CS is dominated by pangenome variants at least one CS in which pangenome variants contributed moderately (0.1 < summed pangenome variant PIP ≤ 0.5), and that CS matched a CS from fine-mapping with 1KG alone, indicating pangenome variants added additional candidates to the signal. **(c)** Example of stable signals, defined as eGenes for which all CSs were matched across both runs and pangenome variants did not contribute meaningfully or not at all (summed pangenome PIP ≤ 0.1), indicating fine-mapping was unaffected by the addition of pangenome-specific variants. **(d)** Example of lost signals, defined as eGenes with no pangenome-refined or pangenome-expanded credible sets, but with at least one CS present in the 1KG alone run that had no matched CS in the merged callset run, indicating that inclusion of pangenome-specific variants diffused previously identified signals.

**Supplementary Figure 9.**
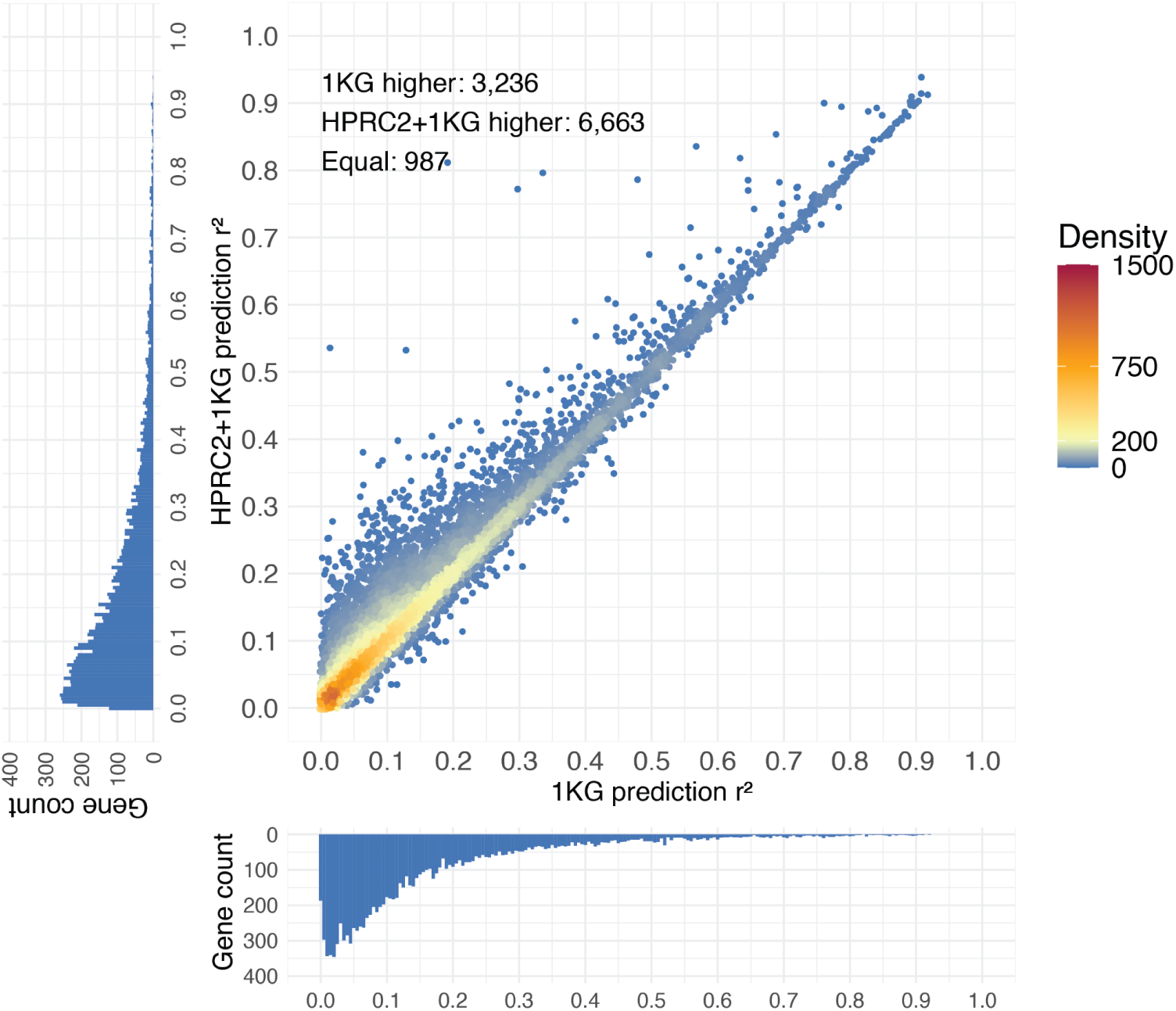
Comparison of gene expression prediction accuracy between the HPRC2+1KG and 1KG callsets. Scatter plot comparing gene expression prediction accuracy for the union of eGenes discovered in eQTL analyses using the 1KG callset or the merged HPRC2+1KG callset (N=10,886). Prediction accuracy was quantified by the 5-fold cross-validated r^2^ between predicted and observed expression, taking the highest adjusted r² across five methods (Lasso, ridge, elastic net, top-1 single best variant, and SuSiE) for each gene within each callset. For each point (eGene), the x axis shows the r^2^ from the 1KG callset, and the y axis shows the r^2^ from the merged HPRC2+1KG callset. Points are colored by local density. Marginal histograms show the distributions of r^2^ for each axis.

**Supplementary Figure 10-14.**
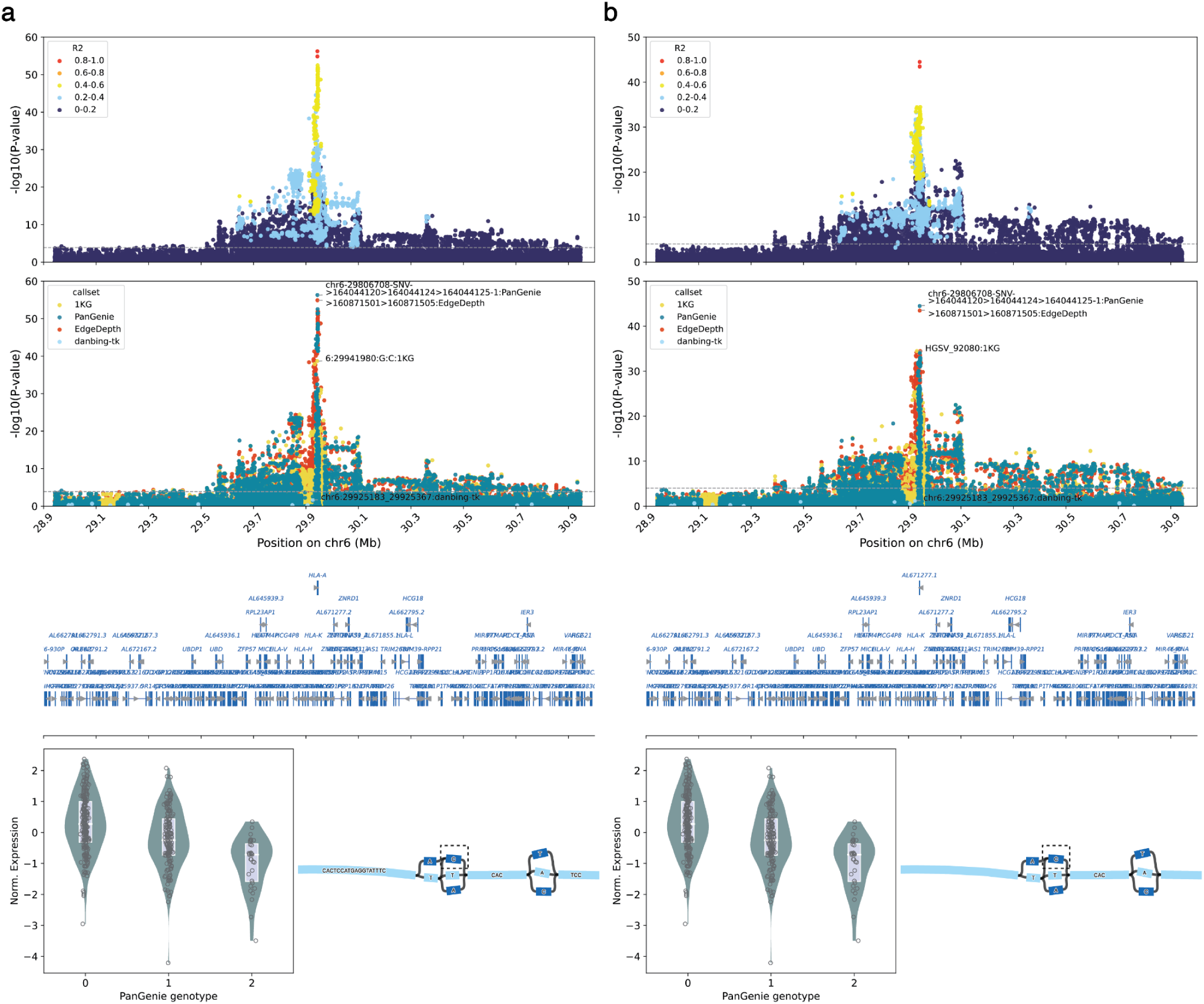
Nine loci showing colocalization of GWAS and eQTL signals. These 9 loci are in addition to the *GBAP1* example shown in Fig. 5, namely *HLA-A* (Supplementary Fig. 10a), *HCG4P5* (Supplementary Fig. 10b), *CREM* (Supplementary Fig. 11a), *EHMT2* (Supplementary Fig. 11b), *UBE2R2* (Supplementary Fig. 12a), *PPM1G* (Supplementary Fig. 12b), *HSD17B8* (Supplementary Fig. 13a), *HCP5* (Supplementary Fig. 13b), and *HMG20A* (Supplementary Fig. 14). Each locus is shown in five panels from top to bottom (as in Supplementary Fig. 3): (1) a LocusZoom plot of association within 1 Mb of the transcription start site (TSS) of the gene color by r^2^ (LD) to the lead eQTL marker; y-axis, −log10(nominal p-value); x-axis, GRCh38 position; the dashed line marks the gene-specific nominal p-value significance threshold; (2) a LocusZoom plot as in (1) colored by callset, with the lead marker of each callset annotated; (3) gene annotations; (4) normalized gene expression versus the lead-marker genotype across 430 AFGR samples. Edge depth genotype is called from fitting a constrained Gaussian mixture model (GMM) from genomeSTRiP to continuous edge depth dosage. Boxes show the median and interquartile range (25th - 75th percentile). Individual samples are shown as points; and (5) a Bandage plot of the pangenome graph structure surrounding the lead variant in the pangenome graph. Reference nodes are colored in light blue and non-reference nodes in dark blue. The lead variant is indicated by an arrow if it is an EdgeDepth variant or by a dashed box if it is a PanGenie variant.

**Supplementary Figure 11.**
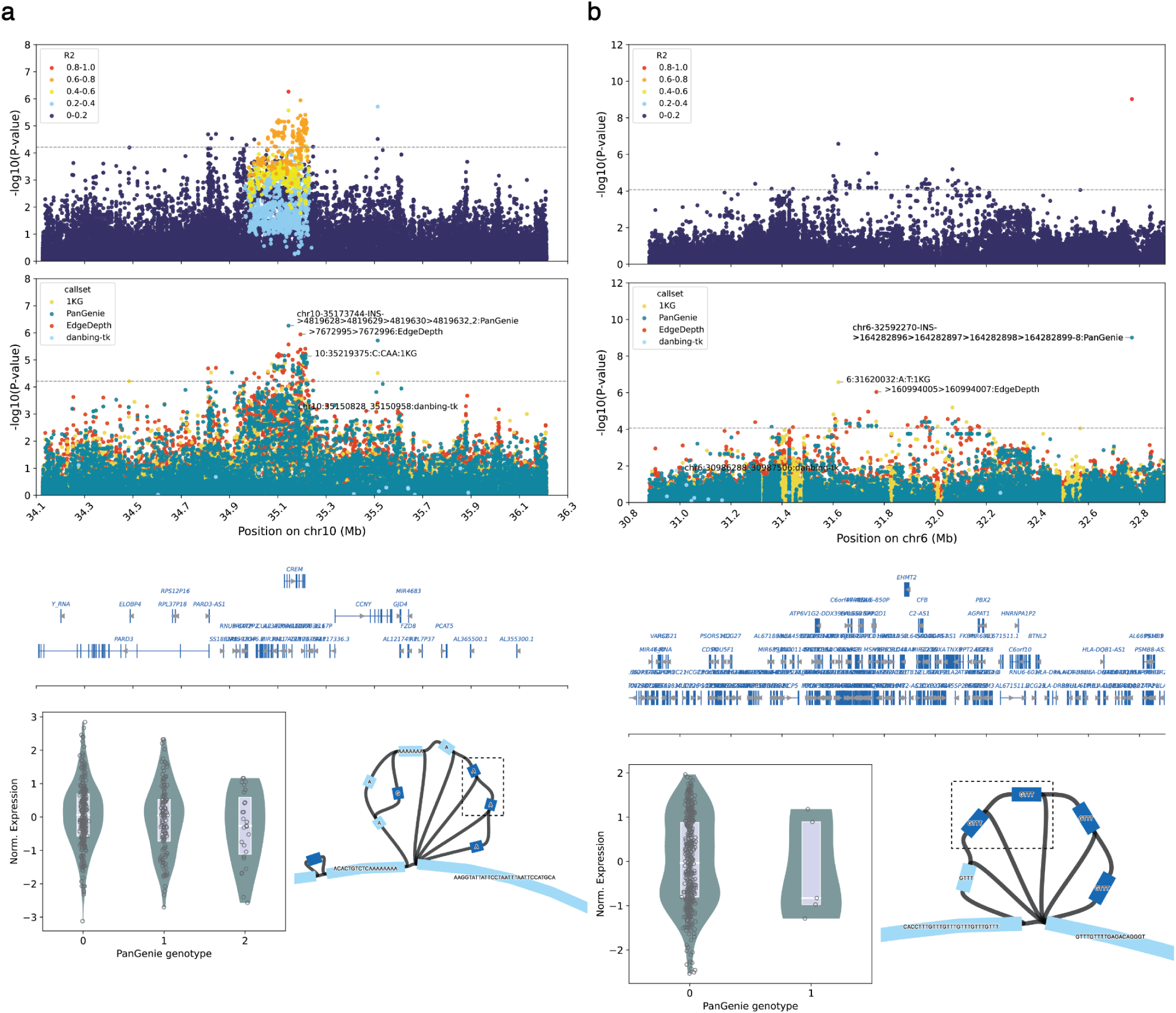
See the legend for Supplementary Figure 10.

**Supplementary Figure 12.**
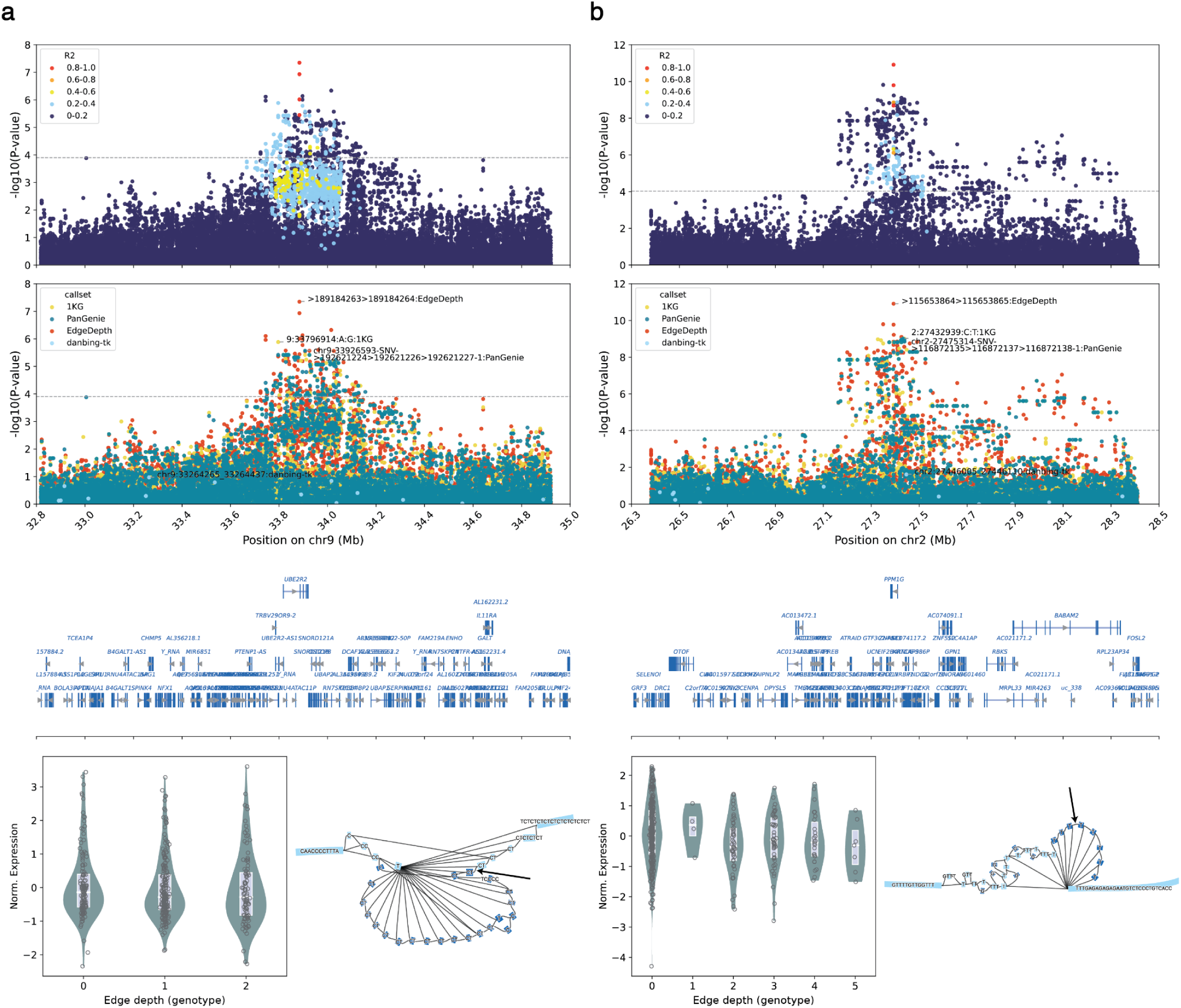
See the legend for Supplementary Figure 10.

**Supplementary Figure 13.**
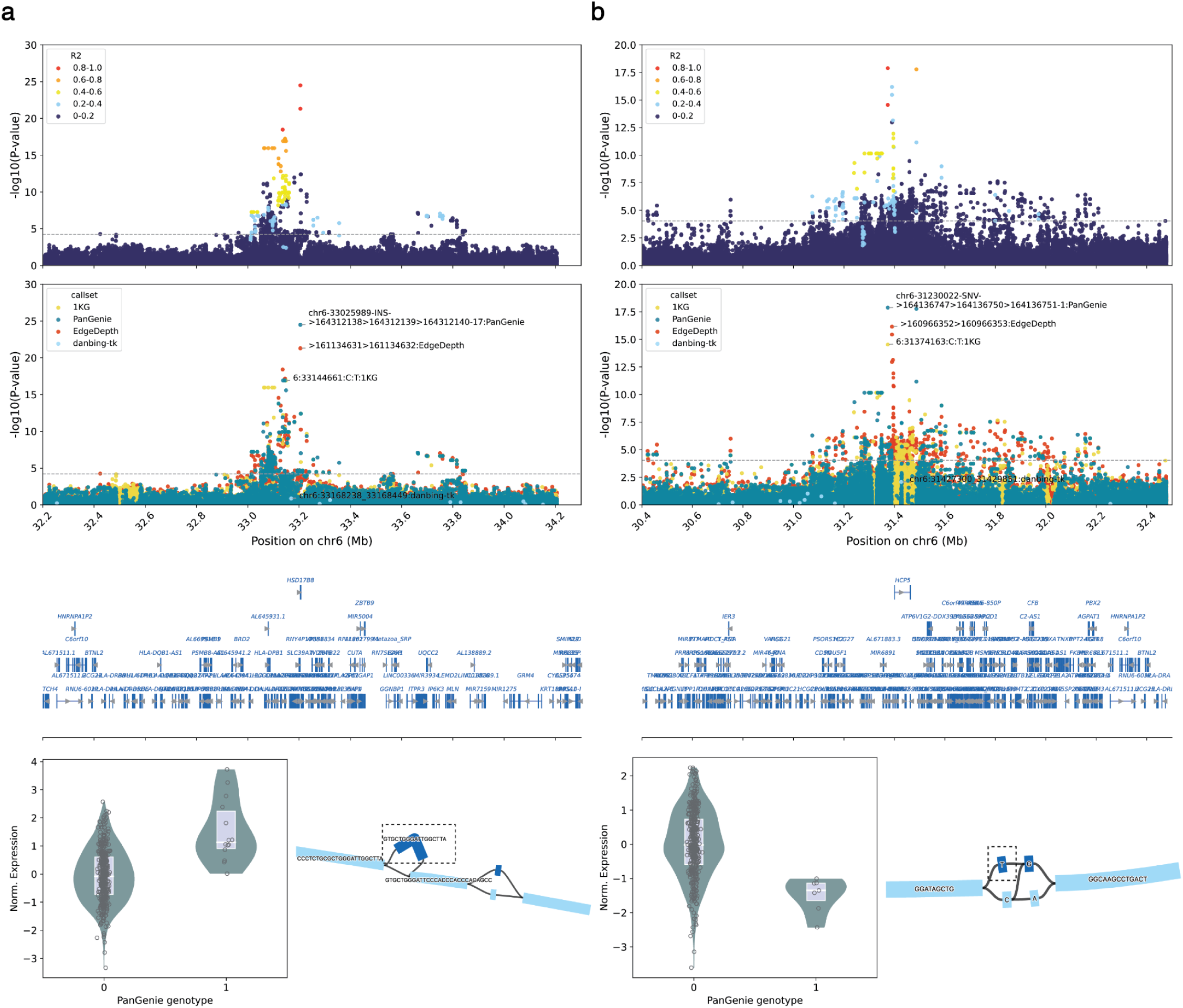
See the legend for Supplementary Figure 10.

**Supplementary Figure 14.**
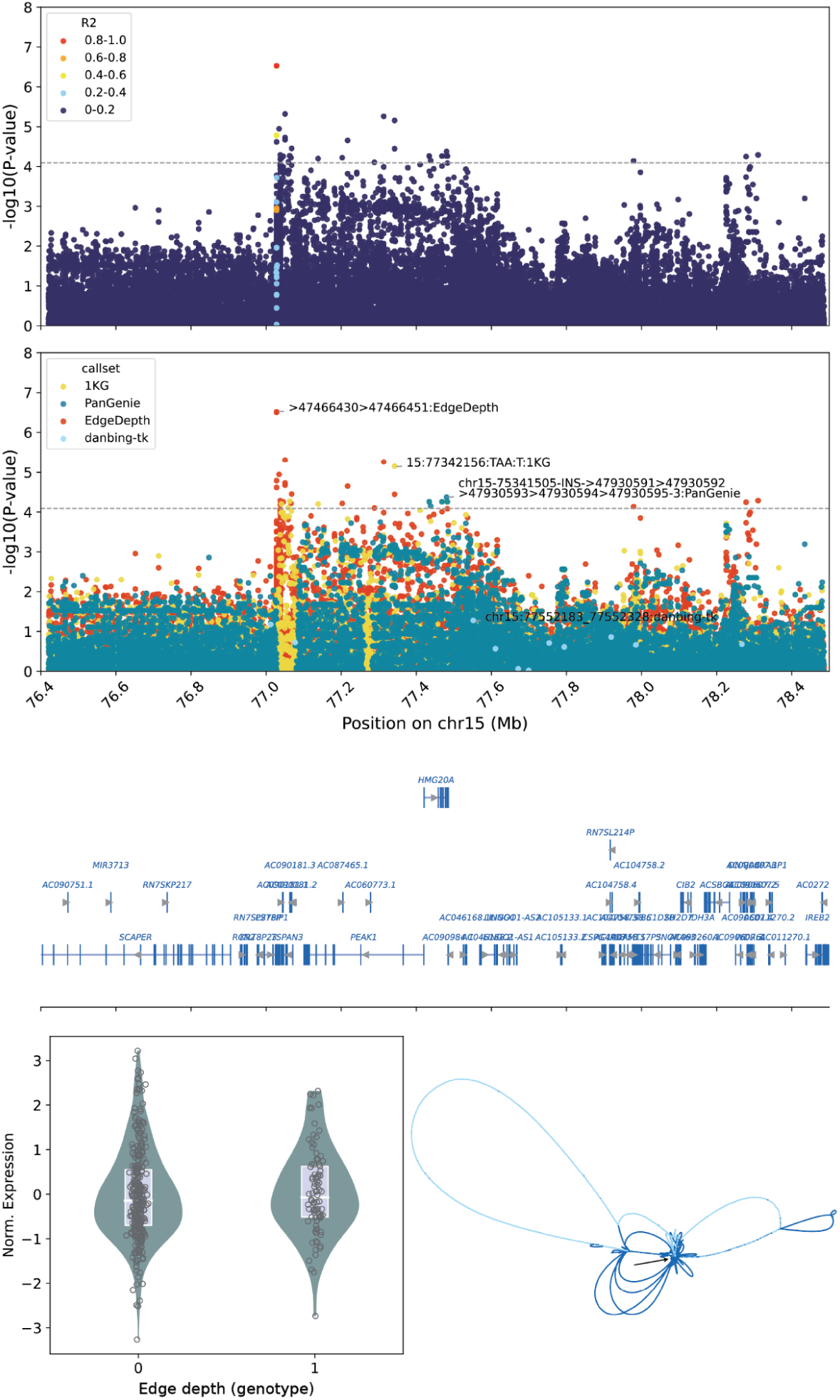
See the legend for Supplementary Figure 10.

**Supplementary figure 15.**
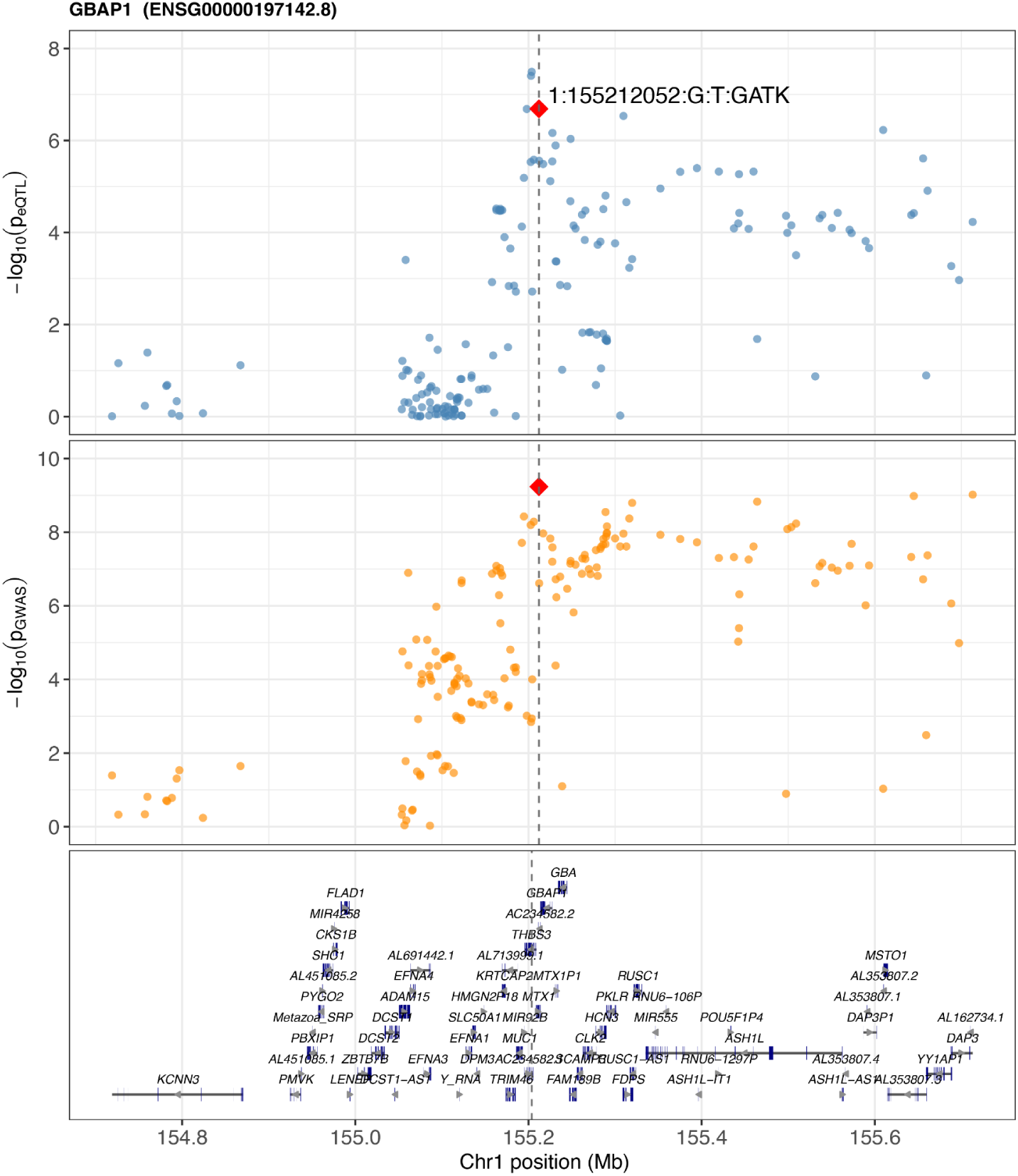
GWAS and eQTL colocalization at the *GBAP1* locus. LocusZoom plot of the *GBAP1* locus showing 1KG variants shared between the eQTL and GWAS analysis within 500 kb of the lead eQTL marker. The top panel shows the eQTL signal at *GBAP1*. The middle panel shows the Crohn’s disease GWAS signal. The bottom panel shows gene annotations in the window. The red diamond marks the variant proposed as the shared causal variant by coloc^31^.

**Supplementary File 1.** Study information for the 41 GWAS used in the GWAS and eQTL colocalization analysis, including study ID, trait, sample size, ancestry, and publication.

**Supplementary File 2.** Summary of the 10 GWAS and eQTL colocalization loci, reporting the eQTL lead variant, eGene, colocalization posterior probability (H4) and GWAS trait at each locus.

